# Type I Interferon-Driven Monocyte Dysregulation and MAS-associated CD8^+^ T cells During Macrophage Activation Syndrome

**DOI:** 10.64898/2026.05.23.727321

**Authors:** Susan P. Canny, Hannah A. DeBerg, Emma L. Kuan, Nicholas Moss, Griffin Gessay, Ailing Lu, Amanda Huang, Allison R. O’Rourke, Erik D. Layton, Paige Bouvatte, Peter J. Wittig, Cate Speake, Carmen Mikacenic, Susan Shenoi, Joyce Hui-Yuen, Daniel J. Campbell, Betsy J. Barnes, Jessica A. Hamerman

## Abstract

Macrophage activation syndrome (MAS) is driven by a hyperinflammatory response characterized by aberrant activation of lymphocytes and phagocytes. While monocytes and macrophages are thought to be important in MAS pathogenesis, their role remains poorly understood. We used bulk and single-cell RNA sequencing (RNA-Seq) on sorted monocytes from children with MAS and healthy controls to identify transcriptional changes during MAS. We defined a MAS signature in classical monocytes that correlated with ferritin and was elevated in monocytes from systemic lupus erythematosus and COVID-19 patients. We also identified a subset of classical monocytes with high levels of interferon-stimulated genes (ISGs) that expanded during MAS. Surprisingly, the transcriptional signature of these cells was driven by type I IFNs, rather than IFNγ. Consistent with this finding, we detected increased levels of circulating IFNβ during MAS, suggesting that IFNβ plays an unrecognized role in driving MAS monocyte responses. We also identified a MAS-associated CD8^+^ T cell population with a distinctive transcriptional signature. We used cell-cell communication algorithms to predict increased immunoregulatory interactions between monocytes and T cells during MAS. Together, these results provide new evidence for a role for type I IFN during MAS and identify a unique CD8^+^ T cell population that may contribute to MAS pathophysiology.

## Introduction

Macrophage activation syndrome (MAS) is a potentially fatal complication of rheumatic diseases, most commonly systemic juvenile idiopathic arthritis (sJIA). sJIA, formerly known as Still’s disease, is an autoinflammatory disorder of unknown etiology characterized by fevers, evanescent rashes, and signs of systemic inflammation accompanied by arthritis or arthralgias. In adults, it is known as Adult-onset Still’s disease (AOSD). MAS affects at least 10% of sJIA patients with up to 50% exhibiting signs of subclinical MAS (1–4). Thus, MAS is an important clinical issue. MAS is identified by the presence of persistent fevers in a patient with known or suspected sJIA/AOSD and specific laboratory findings that are driven by hyperinflammation, including hyperferritinemia (5). While sJIA is typically treated with medications that block the inflammatory cytokines, interleukins 1 and 6 (IL-1 and IL-6), these medications do not alter a patient’s risk of developing MAS (3, 6, 7), implicating other signaling pathways in MAS etiology. Both interferon gamma (IFNγ) and IL-18 have been implicated in MAS (3, 8, 9) and neutralization of IFNγ using an anti-IFNγ antibody, emapalumab, has been effective at inducing MAS remission in a clinical trial (10). However, not all individuals respond to emapalumab, and a better understanding of the immune mechanisms underlying MAS has the potential to identify additional targets for therapeutic interventions.

MAS is a form of secondary hemophagocytic lymphohistiocytosis (sHLH) and shares cellular mechanisms with other forms of HLH. Primary HLH, which is the best understood type of HLH, is typically caused by mutations in genes that contribute to the cytotoxic machinery of CD8^+^ T cells and natural killer (NK) cells. In primary HLH, the lack of CD8^+^ T cell and NK cell cytotoxicity during viral infection leads to a lack of viral clearance and persistent IFNγ production by CD8^+^ T cells and NK cells, driving inflammation that is thought to include overactivation of macrophages (6, 11). However, while a similar inflammatory loop between T/NK cells and macrophages is proposed in MAS, the primary drivers are less clear with a suspected principal role for monocyte dysregulation. Several important studies of monocytes in sJIA have shed light onto their phenotype and function (12–16), and some recent studies have revealed new insights into transcriptional changes in monocytes during MAS (12, 17, 18). Yet, the transcriptional programs that are altered during MAS and how these programs relate to those observed in sJIA remain largely unknown. Circulating monocytes respond both directly and indirectly to the inflammatory milieu and enter tissues where they can differentiate into pathogenic macrophage populations. Therefore, understanding phenotype and function of circulating monocytes during disease states, such as during MAS, can give unique insights into inflammatory diseases.

Previous studies identified emergence of CD38^+^ HLA-DR^+^ CD8^+^ T and NK cells during MAS which distinguish these samples from healthy controls (HC) and sJIA (18–20). These CD38^+^ HLA-DR^+^ lymphocytes, which have also been identified in other forms of HLH, can produce IFNγ, correlate with laboratory disease markers, like CXCL9, IL-18, and ferritin, and are driven by exposure to IFNα/β and IL-15 (18–20). Whether these cells affect monocyte and macrophage activation during MAS has not been assessed.

To address these knowledge gaps, we determined the transcriptomic landscape of monocytes and lymphocytes in sJIA with and without MAS compared to HCs using bulk and single-cell RNA sequencing (RNA-Seq) and cell-cell interaction analyses. Our monocyte analyses revealed that classical monocytes during MAS demonstrated a type I interferon (IFNα/β) signature whereas nonclassical monocytes were the only population to show an IFNγ signature during MAS. We also directly detected elevated levels of circulating IFNβ in all MAS participants. In addition, we identified a MAS-associated CD8^+^ T cell population with a unique phenotype defined by expression of the proliferative marker *CD27*, the homing receptor *CXCR6*, and multiple inhibitory receptors. Cell-cell interaction analyses uncovered immunoregulatory interactions between monocytes and T cells. Together, our studies revealed an important role for IFNβ during MAS, identified a population of MAS-associated CD8^+^ T cells, and suggested that immunoregulatory pathways were activated during MAS.

## Results

### Study design and participant characteristics

To better understand how MAS effects circulating immune cells, we collected peripheral blood mononuclear cells (PBMCs) and plasma from individuals presenting with sJIA with MAS (MAS), sJIA without MAS (active sJIA), or quiescent sJIA and no MAS (quiescent) for analysis using bulk and single-cell RNA-Seq and measurement of circulating cytokines (Supplemental Figure 1). PBMCs and plasma from HCs were selected from samples available in the Benaroya Research Institute biorepository. Individuals with MAS were identified based on the Ravelli criteria for MAS (5). sJIA was diagnosed by the treating pediatric rheumatologist. Quiescent disease was determined by the treating pediatric rheumatologist and subjects were without fever, rash, arthritis, or notable laboratory abnormalities. Participants had a mixture of treatments (Supplemental Tables 2-5) at the time of sample collection with some individuals on no medications and others on a mix of immunomodulatory treatments. For all RNA-Seq experiments, and all but one plasma sample, MAS samples were collected prior to the initiation of systemic steroids. For those individuals on medications at the time of sample collection, length of treatment varied widely as some individuals were newly diagnosed and others had a disease flare while on medications and had been diagnosed months to years prior.

### Identification of a MAS gene signature in classical monocytes

Because macrophage inflammatory responses are thought to be involved in MAS, we examined circulating monocytes as they are an accessible population of cells in patients that have the capacity to differentiate into macrophages and are key mediators of inflammatory responses. We performed bulk RNA-Seq on sorted classical CD14^+^CD16^-/low^ monocytes from cryopreserved PBMCs from individuals with MAS (n=6), active sJIA (n=4), quiescent disease (n=8), and HCs (n=8) (Supplemental Figure 2A, Supplemental Tables 1-2). During sorting, we found that the major population of CD14^+^ monocytes from MAS individuals had significantly higher CD16 expression on their cell surface compared to those from active sJIA or HCs (Figure 1A), thus we sorted this entire population (Supplemental Figure 2B). This increased CD16 expression appeared distinct from CD14^+^CD16^+^ intermediate monocytes (Supplemental Figure 2B and data not shown). However, after three days of high-dose methylprednisolone treatment, the CD16 expression on classical monocytes decreased to a level comparable to that seen for HCs (Supplemental Figure 3A). In contrast, the surface expression of CD163, a hemoglobin-haptoglobin scavenger receptor, was increased on MAS monocytes after initiation of steroid treatment (Supplemental Figure 3B); CD163 is cleaved from the cell surface upon cell activation, and soluble CD163 in serum can be used as a marker of MAS (21, 22). Therefore, the reduction in CD16 expression and increase in CD163 expression after steroid treatment most likely reflects a reduction in overall monocyte activation during treatment of MAS.

**Figure 1:**
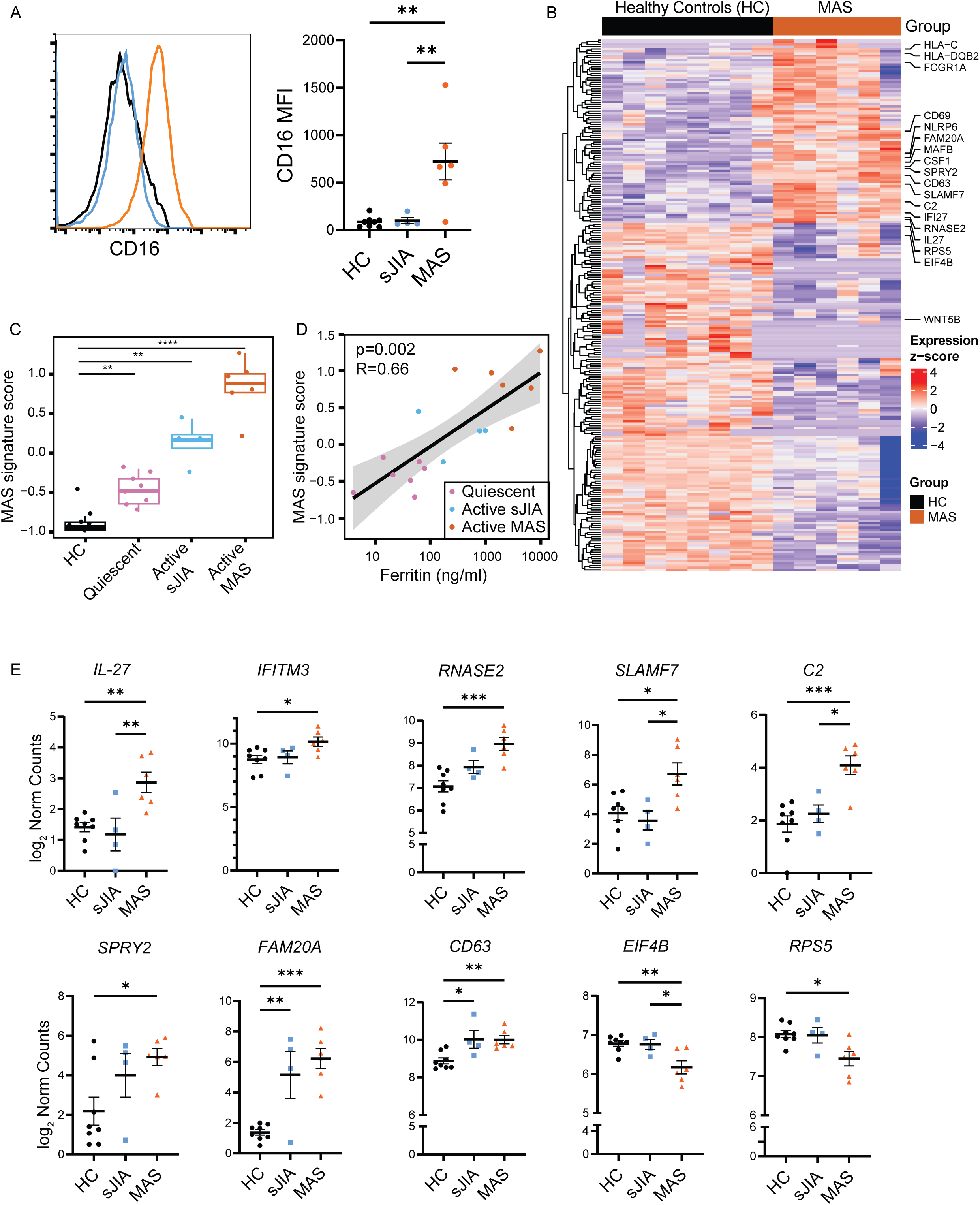
Definition of MAS monocyte signature using bulk RNA-Seq. (A) PBMCs from HC (n=8), sJIA (n=4) or MAS (n=6) were analyzed using flow cytometry. Monocytes were gated as CD3^-^CD19^-^CD56^-^CD15^-^CD14^high^HLA-DR^+^ cells and the expression of CD16 was determined. Histogram (left) shows representative sample and graph (right) shows mean fluorescent intensity (MFI) for each sample. (B) Heatmap showing DEGs between MAS and HC, defined as genes with FDR < 0.1, from sorted CD14^+^CD16^-/low^ monocytes (sorting scheme shown in Supplemental Figure 1A). Z-scores for DEGs are displayed for individual participants (columns). (C) MAS signature score was generated from DEG in (B), and then applied to data from all individuals, including HC (black), quiescent disease (pink), active sJIA without MAS (blue) and active MAS (orange). (D) MAS signature scores were positively correlated with serum ferritin levels from all disease samples in (C). (E) Normalized RNA read counts for specific transcripts of interest. Mean +/- SEM, each dot represents the values for an individual sample (A, C, E). * < 0.05, ** < 0.01, *** < 0.001 by one way ANOVA with Tukey’s multiple comparisons test. For RNA-Seq data (panels B-E), HC (n=8), quiescent (n=8), sJIA (n=4), and MAS (n=6).

We identified 240 differentially expressed genes (DEGs) that distinguished individuals with MAS from HCs with an FDR < 0.1 (Figure 1B, Supplemental Data 1). At an FDR < 0.05, there were 95 DEGs that distinguished MAS from HC; we used these 95 DEGs to generate a MAS signature score in classical monocytes, defined as the difference between mean expression of upregulated and downregulated DEGs. As expected, this signature was highest in classical monocytes from individuals with MAS (Figure 1C). We also applied this signature to classical monocytes from children with active sJIA or with quiescent disease (Supplemental Figure 3A). We observed a stepwise increase in the MAS signature across groups with HCs < quiescent disease < active sJIA < MAS (Figure 1C). For the two individuals with MAS who had longitudinal samples obtained before and after three days of steroid treatment, the MAS signature score was substantially reduced following treatment (Supplemental Figure 2C), consistent with improvement in clinical status and lab values. We also found that the classical monocyte MAS signature score positively correlated with ferritin, a laboratory test used in clinical practice to identify those with MAS (Figure 1D).

Although pathway enrichment analysis did not identify any pathways significantly enriched in the MAS gene signature, a review of individual DEGs upregulated in classical monocytes from individuals with MAS revealed several genes of interest (Figure 1E). For example, some transcripts were specifically upregulated during MAS but not active sJIA when compared to HC, such as *IL-27*, *IFITM3*, *RNASE2*, *SLAMF7*, and *C2* (Figure 1E). Interestingly, *IL-27*, *IFITM3*, and *RNASE2* can all be induced by IFNα/β. In contrast, other transcripts such as *SPRY2*, *FAM20A*, and *CD63*, were upregulated in both MAS and active sJIA monocytes (Figure 1E) (23). There were also many genes that were downregulated specifically during MAS, including *EIF4B* and *RPS5*, which are involved in protein translation (Figure 1E).

We next used our monocyte MAS signature score to analyze publicly available datasets. We first applied our score to an independent monocyte dataset from sJIA participants (12) and found that new onset sJIA and MAS individuals had higher MAS signature scores whereas active sJIA, quiescent disease, and HCs had a lower MAS signature score (Supplemental Figure 4B, (12)). Consistent with our data, there was also a significant correlation between ferritin and the MAS monocyte signature score in this dataset (Supplemental Figure 4B, (12)). To understand whether this score has utility in other systemic inflammatory diseases, we compared the MAS monocyte signature score to publicly available data from childhood systemic lupus erythematosus (cSLE) (24). We identified an increased MAS signature score in classical monocytes from cSLE (Supplemental Figure 4C) and found a correlation between the SLE disease activity index (SLEDAI) and MAS signature score (Supplemental Figure 4C), suggesting that there are shared transcriptional changes in monocytes during active SLE and active MAS. Next, we tested whether this signature score was elevated in a distinct setting of strong systemic inflammation as seen during viral infection with SARS-CoV2. We analyzed classical monocytes sorted from cryopreserved PBMCs from adults hospitalized with COVID-19 in 2020 and 2021 using bulk RNA-Seq (25). We found that our MAS monocyte signature score was also increased during this severe viral infection (Supplemental Fig 4D). Taken together, these data define a monocyte MAS signature score strongly correlated with ferritin in sJIA-driven MAS and that captured a shared inflammatory program also present in SLE and serious SARS-CoV2 infection.

### scRNA-sequencing identifies eight myeloid clusters in HC and MAS

To better characterize the diversity of monocyte responses during MAS, we performed single-cell RNA-Seq (scRNA-Seq) on sorted total monocytes from PBMCs from individuals with MAS (n=8) or from HCs (n=8; Supplemental Figure 5, Supplemental Tables 1 and 3). All MAS samples were collected prior to treatment with systemic corticosteroids (Supplemental Table 3). When we performed clustering to identify monocyte subsets, we noted a strong effect of individual donors in classical monocyte clusters that was not seen in nonclassical monocytes or in the few plasmacytoid dendritic cells (pDCs) in these samples (data not shown). We computationally removed donor-specific effects observed in the classical monocyte clustering using integration with the R package Seurat (26). Applying reference labeling using the Human Immune Healthy Atlas (27), we found that, as expected, most of the monocytes were identified as CD14^+^ monocytes with a smaller population of CD16^+^ monocytes (Figure 2A). Unbiased clustering revealed seven CD14^+^ clusters and one cluster, Cluster 4, with low *CD14* expression and high *FCGR3A*, encoding CD16, along with complement components, such as *C1QA*, consistent with nonclassical monocytes (Figure 2B-D).

**Figure 2:**
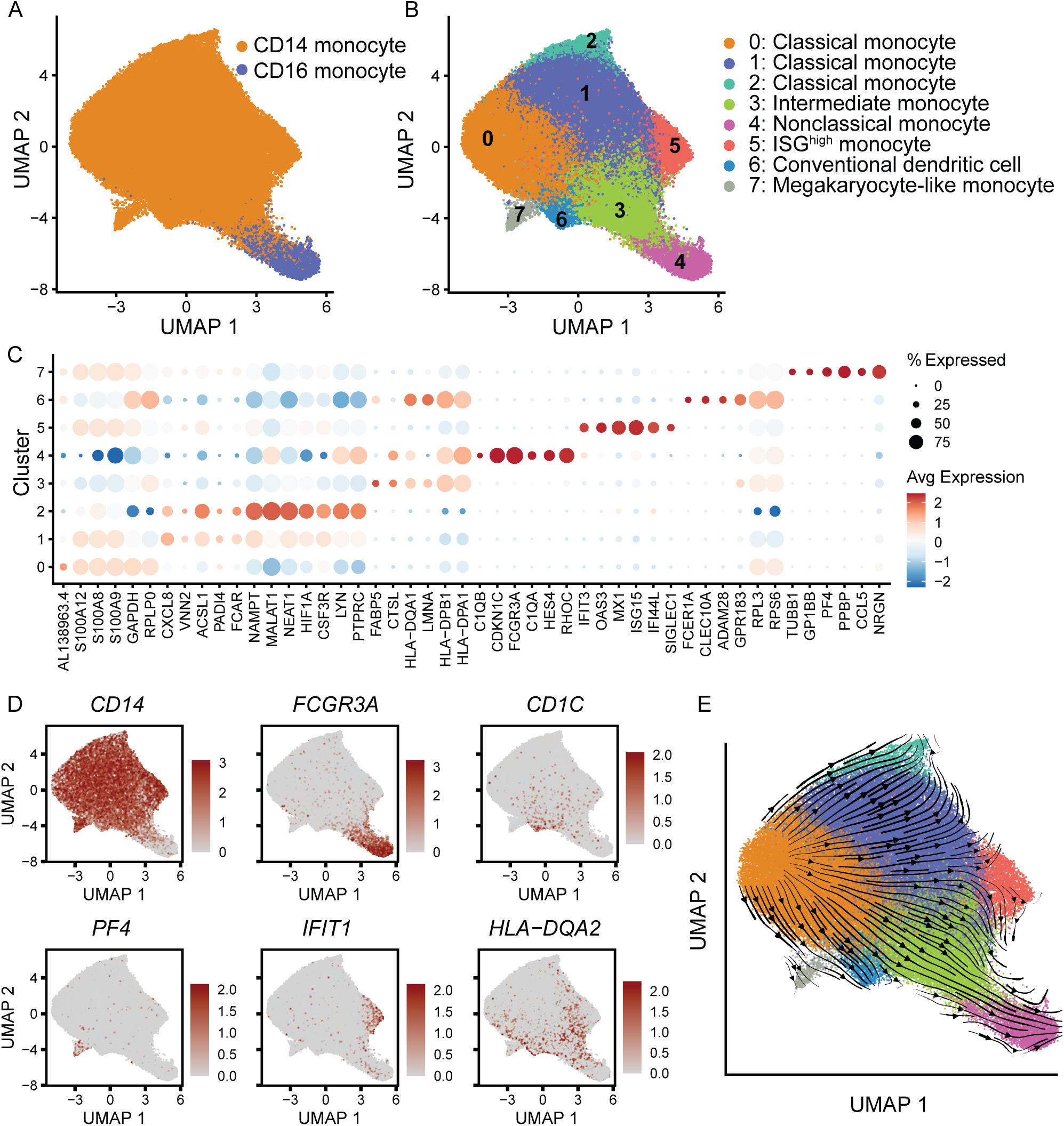
Defining the MAS monocyte landscape using scRNA-Seq. (A) UMAP with cell type annotations from mapping to a published dataset (27). (B) UMAP with unsupervised clustering of monocytes identified eight monocyte clusters. (C) Dot plot showing expression of selected cluster-defining genes chosen from among the top 20 DEGs per cluster. Dot size is proportional to the percentage of cells expressing the gene in each cluster and color reflects the average scaled log-normalized expression within a cluster. (D) Expression of selected cluster-defining genes. (E) RNA velocity trajectory inference (arrows) overlaid on the monocyte UMAP from (B).

We further classified the clusters of CD14^+^ monocytes by examining gene expression and cluster defining genes (Supplemental Data 2). Cluster 3 had increased expression of class II HLA molecules, consistent with intermediate (CD14^+^CD16^+^) monocytes (Figure 2B-D). We also identified a small cluster of conventional dendritic cells (cDCs, Cluster 6) that expressed *CD1C* and *CLEC10A* transcripts (Figure 2B-D). Cluster 7 consisted of *CD14^+^* cells with high levels of transcripts typically associated with platelets including *PF4*, *GP9*, and *PPBP* (Figure 2B-D).

Similar clusters of “megakaryocyte-like” monocytes have been reported in other scRNA-Seq datasets (28) and are likely doublets of platelets and monocytes but could also represent expression of platelet genes in monocytes due to differentiation state or phagocytosis.

At this resolution, we identified four classical monocyte clusters (Clusters 0, 1, 2, and 5; Figure 2B). Clusters 0, 1, and 2 exhibited subtle differences between each other (Figure 2C).

While all classical monocytes expressed the alarmins *S100A8*, *S100A9*, *S100A12*, these were cluster-defining genes for Cluster 0 (Figure 2C). Multiple ribosomal genes were also cluster-defining genes for Cluster 0. Cluster 1 defining genes included the neutrophil-attracting chemokine *CXCL8,* whereas Cluster 2 defining genes included increased expression of long noncoding RNAs, *MALAT1* and *NEAT1*, and downregulation of multiple ribosome genes, such as *RPL0* (Figure 2C). Cluster 5 was notable for IFN-related cluster-defining genes including *IFIT3, OAS3, MX1, ISG15, IFI444L,* and *SIGLEC1* (Figure 2C). This population of high interferon-stimulating genes (ISG^high^) monocytes has been previously identified by others in both HCs and inflammatory diseases and is expanded in SLE (24, 27, 29).

To investigate relationships between monocyte clusters, we performed RNA velocity analysis, which infers cell differentiation trajectories using the ratio of unspliced to spliced transcripts from scRNA-Seq data (30). RNA velocity predicted the known differentiation pathway from classical monocytes (Cluster 0) to intermediate monocytes (Cluster 3) to nonclassical monocytes (Cluster 4; Figure 2E), thus validating this approach. RNA velocity also indicated differentiation of Cluster 0 into other subsets of classical monocytes (Clusters 1 and 2), suggesting that classical monocyte Cluster 0 may represent the CD14^+^ monocytes that enter circulation from the bone marrow. Interestingly, RNA velocity predicted minimal differentiation of any monocyte cluster into the ISG^high^ monocytes (Cluster 5; Figure 2E). This finding suggests that ISG^high^ monocytes may not differentiate from other monocytes in circulation, but instead derive directly from a distinct bone marrow progenitor or differentiate in peripheral tissues rather than from circulating monocytes. Alternatively, gene expression in ISG^high^ monocytes may be at a transcriptional steady-state.

### The transcriptome of MAS monocytes is distinct from healthy monocytes

To assess how MAS and HC monocytes differ, we first compared the proportion of MAS to HC monocytes per cluster. All clusters contained cells from both MAS and HC subjects (Figure 3A). However, only the ISG^high^ monocyte cluster (Cluster 5) had significant differences between MAS and HC with a higher percentage of MAS cells in this cluster (Figure 3A and Supplemental Figure 5A). There was also a trend for a lower percentage of the nonclassical monocyte cluster in MAS (Cluster 4; Figure 3A). We next asked whether our MAS signature score was specific to certain types of monocytes by applying it to pseudobulked monocyte clusters observed in the single-cell data. We found that the MAS monocyte gene signature was upregulated in all eight MAS myeloid clusters compared to HC and was most pronounced in the classical and intermediate clusters (Supplemental Figure 6B), indicating global enrichment of this signature. This suggests that a shared inflammatory environment shapes similar responses in all circulating monocytes during MAS.

**Figure 3:**
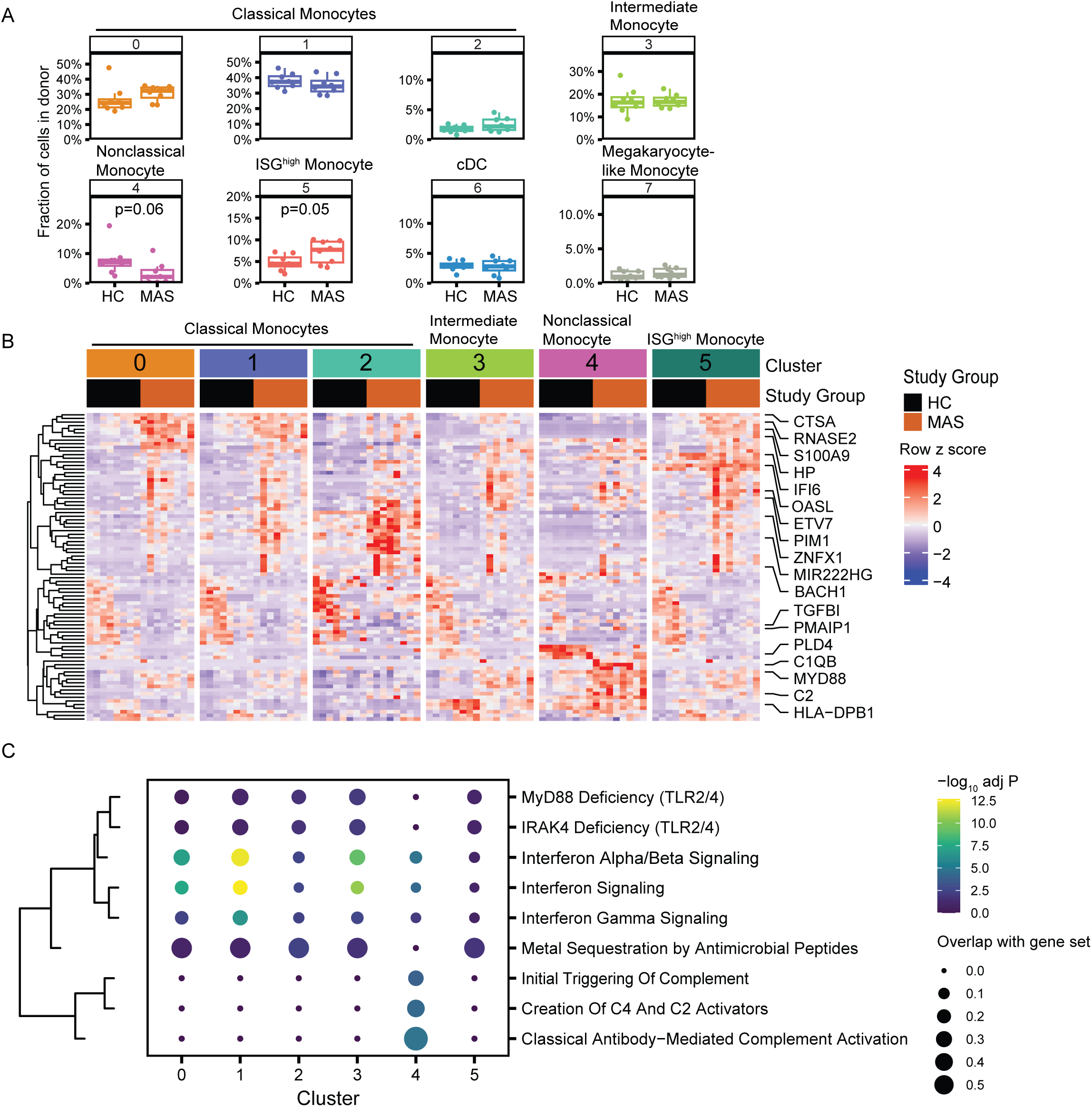
MAS-specific changes in monocytes by scRNA-Seq. (A) Cluster proportions between HC and MAS participants, each dot represents an individual. p-values were calculated using Student’s unpaired t-test. (B) DEGs across major monocyte clusters. Gene expression data was pseudobulked to the level of donor and cluster. The top 20 genes in each cluster with FDR < 0.05 and a log2-fold change > 0.5 for any cluster are shown. (C) Reactome enrichment of pseudobulk MAS signatures on the genes up in MAS relative to HC within each cluster. Dot size indicates overlap with the Reactome gene set and color represents enrichment p-value.

To define DEGs between MAS and HC in each cluster, we used pseudobulk analysis; we identified DEGs that were shared across clusters and others that were specific to individual clusters (Figure 3B, Supplemental Data 3). For example, the pro-survival molecule, *PIM1*, was increased in all clusters of MAS monocytes compared to HC monocytes whereas *PMAIP1*, which encodes a proapoptotic protein, was decreased in all MAS monocyte clusters (Figure 3B). These shared gene expression changes may contribute to prolonged survival of MAS monocytes. *TGFBI*, a transcript induced by the immunoregulatory cytokine TGFβ, was also decreased in all MAS monocyte clusters. Some genes were differentially expressed specifically in classical MAS monocytes (Clusters 0-2, 5); these included higher expression of genes that are commonly expressed in classical monocytes, such as the alarmin, *S100A9*, and cathepsin A, *CTSA* (Figure 3B). In addition, *HP,* which encodes haptoglobin and binds free hemoglobin, was also more highly expressed in MAS classical monocytes (Clusters 0-2 and 5)C. ISGs, such as *IFI6* and *OASL,* were also increased in MAS cells from most monocyte clusters (Clusters 0-4); notably, ISGs were highly expressed in ISG^high^ monocytes (Cluster 5) from both MAS and HC (Figure 3B).

Some DEGs had particularly strong expression in MAS cells from specific clusters. For example, genes associated with viral RNA sensing, such as *ZNFX1* and *TRIM25*, and *MIR222HG*, which is involved in monocyte/macrophage polarization, were strongly upregulated in MAS cells from Cluster 2 compared to the other classical monocytes (Clusters 0, 1, and 5).

We also identified a set of DEGs that were primarily differentially expressed in nonclassical MAS monocytes (Cluster 4), including decreases in *SEZ6L*, involved in complement regulation, and *PLD4*, involved in phagocytosis and regulation of the inflammatory response. There was a larger set of genes that were primarily increased in MAS nonclassical monocytes which included complement components, *C1QB* and *C2*, in addition to the toll-like receptor (TLR) signaling adaptor, *MYD88*. In sum, these observations reveal both shared and cluster-specific MAS-induced transcriptional changes.

Pathway analysis of DEGs (Figure 3B) found that IFNα/β and IFNγ signaling pathways were enriched across MAS monocyte clusters, especially in classical monocytes (Clusters 0 and 1) and the intermediate monocytes (Cluster 3; Figure 3C). IFN signaling in the ISG^high^ cluster (Cluster 5) was likely less enhanced in MAS compared to HC monocytes due to baseline expression of ISGs in HC ISG^high^ cells. Complement pathways were particularly enriched in MAS nonclassical monocytes (Cluster 4), while the Metal Sequestration by Antimicrobial Peptides pathway was enriched in all other MAS monocyte clusters (Figure 3C). This metal sequestration pathway includes the alarmins, *S100A7*, *S100A8*, and *S100A9*, which are during active sJIA (31, 32). In summary, pathway analysis highlighted the importance of IFN signaling across all monocytes as well as distinct ways nonclassical and classical monocytes responded during MAS.

### Type I IFN signature in MAS monocytes

Although IFNγ plays an important role in MAS and other forms of HLH (6, 10, 33–35), a role for IFNα/β is less explored. However, we noted several of the ISGs contributing to our MAS signature are more often associated with IFNα/β signaling than IFNγ signaling. Using a publicly available cytokine dictionary generated in mice treated with cytokines in vivo (36), analysis of our bulk MAS monocyte DEGs using mouse orthologs yielded strong evidence for the presence of an IFNβ response as well as an IFNα1 response (Figure 4A), corroborating data by Huang et al. demonstrating the presence of IFNα/β activity during MAS (18). In contrast, the IFNγ response did not reach statistical significance (Figure 4A). Notably, IL-15 and IL-18 were also significantly enriched in the MAS monocyte signatures (Figure 4A), both of which have been associated with MAS (8, 18, 37). Taken together, these results suggest that IFNα/β is important in shaping monocyte responses and provide further evidence of the importance of IL-15 and IL-18 during MAS.

**Figure 4:**
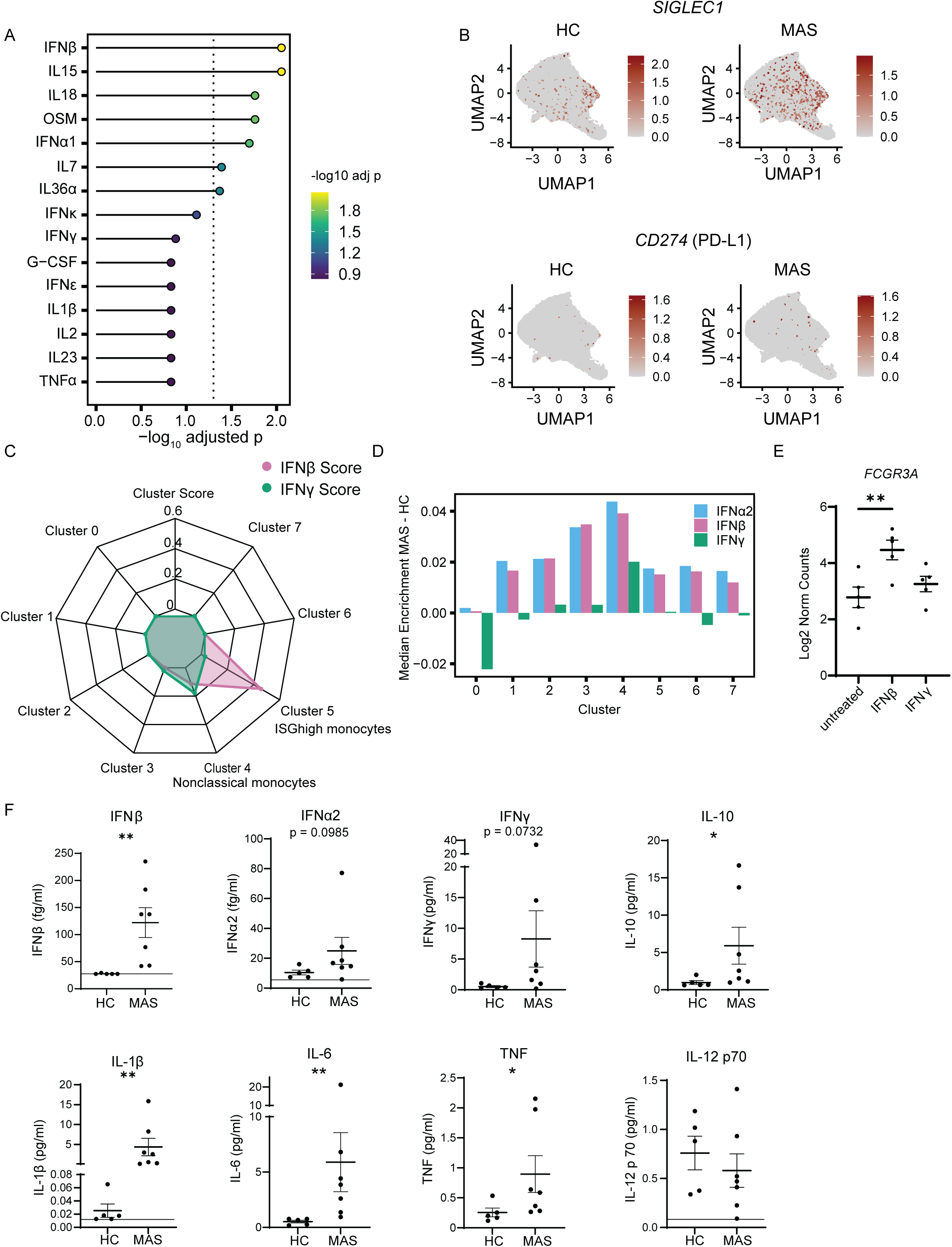
A type I IFN signature in MAS monocytes and circulating IFNβ during MAS. (A) Cytokine signature analysis of genes upregulated (FDR < 0.1) in bulk RNA-Seq MAS CD14^+^ monocyte samples relative to HC (36). Color and x-axis represent enrichment p-values. (B) UMAP colored by expression of *SIGLEC1* and *CD274* in HC (left) or MAS (right) cells; downsampled to 10,000 cells per UMAP. (C) Radar plot showing IFN-β and IFN-γ scores per cluster obtained from non-negative least squares modeling coefficients of monocytes treated with the relevant cytokine. (D) Bar plots showing mean MAS enrichment minus mean HC enrichment from AUCell analysis using IFNα2, IFNβ, and IFNγ signatures. (E) Log_2_-normalized expression of *FCGR3A*, encoding CD16, in bulk RNA-seq of monocytes from healthy participants stimulated with IFNβ, IFNγ, or left untreated (n = 5). One way ANOVA with Tukey’s multiple comparisons test. (F) Plasma concentrations of cytokines in HC and MAS samples. For those samples below the limit of assay detection, the concentration was set to the level of detection (indicated with a horizontal line). For samples below the limit of quantitation but above the limit of detection, the value measured is displayed. Mann Whitney U test. *p < 0.05, **p < 0.01 (E, F).

To further dissect the relative contributions of IFNα/β and IFNγ in MAS monocyte gene expression, we assessed the expression of *SIGLEC1* as an IFNα/β-induced gene and *CD274* as an IFNγ-induced gene (38–40), because expression of these proteins by flow cytometry is used as a clinical assay to distinguish IFNα/β vs IFNγ signaling. *SIGLEC1* expression was enriched in the ISG^high^ monocyte cluster (Cluster 5) in HC individuals whereas it was expressed in all clusters except for the nonclassical monocyte cluster (Cluster 4) in MAS individuals (Figure 4B), supporting a broad IFNα/β signature in classical monocytes during MAS. In contrast, *CD274* expression was low overall in both HC and MAS groups, though more MAS monocytes had detectable *CD274* than HC monocytes (Figure 4B). These findings suggest that IFNα/β may play a more prominent role during MAS than previously recognized.

To take a broad view of IFNα/β and IFNγ induced transcriptional changes, we applied IFN gene sets generated from human monocytes incubated with IFNα2, IFNβ, or IFNγ compared to untreated monocytes using scRNA-Seq (41). A non-negative matrix factorization (NMF) approach using the IFNβ vs IFNγ signatures was used to decompose IFN signaling in our single-cell data into IFNβ or IFNγ components. Of note, genes induced by IFNα2 or IFNβ in human monocytes were nearly identical, precluding inclusion of both IFNα2 and IFNβ in our NMF model. This analysis showed that IFNβ signaling strongly contributed to gene expression in the ISG^high^ monocyte cluster (Cluster 5) with a much weaker IFNγ signature in this population (Figure 4C). In contrast, nonclassical monocytes (Cluster 4) exhibited a stronger IFNγ signature and a weaker IFNα/β signature than ISG^high^ monocytes (Cluster 5; Figure 4C).

To determine the contribution of specific IFNs during MAS, we compared the enrichment of each IFN signature to HC monocytes for each cluster. All monocyte clusters except Cluster 0 had enrichment of the IFNα/β signatures in MAS compared to HC, with the strongest enrichment in nonclassical (Cluster 4) and intermediate monocytes (Cluster 3; Figure 4D, Supplemental Figure 6A). Nonclassical MAS monocytes (Cluster 4) showed the strongest enrichment for the IFNγ signature although this was less pronounced than the IFNα/β signature. No other monocyte clusters showed appreciable enrichment for the IFNγ signature during MAS. Interestingly, the classical monocytes in Cluster 0 had a relatively strong downregulation of the IFNγ signature and little IFNα/β signature during MAS, suggesting they were either refractory to IFNα/β signaling or were not exposed to IFNα/β. Overall, these data demonstrate a key role for broad IFNα/β signaling in monocytes during MAS, with a more focused role of IFNγ signaling in nonclassical monocytes.

Given evidence of an IFNα/β signature in MAS monocytes and the upregulation of CD16 that we observed specifically on the surface of MAS classical CD14^+^ monocytes (Figure 1A), we next asked whether IFNβ or IFNγ induces transcription of *FCGRIIIA*, which encodes CD16. We incubated monocytes from HCs with IFNβ or IFNγ or media alone and then collected RNA for bulk RNA-Seq. We found that IFNβ, but not IFNγ, robustly induced transcription of *FCGRIIIA* (Figure 4E), indicating that IFNβ likely accounts for the upregulation of CD16 we observed on MAS classical monocytes (Figure 1A).

The IFNα/β signature in MAS monocytes led us to investigate whether these cytokines were present in circulation during MAS using a high sensitivity assay to measure IFNβ and IFNα2 in plasma from MAS (n=7) and HC (n=5) individuals (Figure 4F, Supplemental Tables 1, 5). IFNβ was readily detected in all MAS plasma samples and was significantly higher in MAS plasma relative to HC (Figure 4F). IFNα2 protein was above the level of detection in all participants and trended higher in MAS participants (Figure 4F) although this difference was primarily driven by two individuals. Consistent with the known low baseline type I IFNs in circulation, most of the HC plasma samples had undetectable or very low IFNβ or IFNα2. IFNγ was easily detectable in all samples and trended higher in MAS participants (Figure 4F).

In contrast to individuals with MAS, individuals with active sJIA or quiescent disease had low levels of all IFNs measured (Supplemental Figure 7B). While IFNα2 and IFNγ levels were comparable to HCs, the level of IFNβ in active sJIA was modestly higher than HCs but below the level seen in MAS (Supplemental Figure 7B). Of the quiescent samples, the three with highest IFNβ levels had a history of MAS and the highest quiescent IFNβ sample was from a subject with sJIA lung disease. For those MAS samples with a paired quiescent sample, we observed a higher IFNβ level during active MAS (Supplemental Figure 7C). Taken together, these data are consistent with our scRNA-Seq data which show strong evidence of an IFNα/β signature in MAS monocytes across all monocyte clusters.

We measured additional plasma cytokines based on their association with sJIA or MAS. We found that the plasma concentrations for IL-1β, IL-6, TNF, and IL-10, but not IL-12p70, were significantly higher during MAS than in HCs (Figure 4F). Of note, IL-1β and IL-6 concentrations in plasma were higher in active sJIA than in MAS while IFNβ, IFNα2, IFNγ, and IL-10 were higher in MAS than active sJIA (Supplemental Figure 7B), supporting the hypothesis that IFNs and IL-18 drive MAS activity while IL-1 and IL-6 drive sJIA activity (3, 6, 10, 37, 42). In three individuals with paired samples from MAS and quiescent disease, IFNγ and IL-10 were higher during active MAS (Supplemental Figure 7C). Therefore, we found that MAS participants with active disease had specific elevation in IFNβ, IFNγ, and IL-10, while sJIA participants demonstrated increased levels of IL-1β and IL-6.

### Identification of MAS-associated CD8^+^ T cells in the CD8^+^ effector/memory compartment

Given the importance of CD8^+^ T cells and NK cells during HLH, we also performed scRNA-Seq on sorted lymphocytes from a subset of our participants (n=6 HC, n=6 MAS; Supplemental Tables 1, 4, Supplemental Figure 4). Reference mapping using the Immune Cell Atlas (27) identified T, B, and NK cell subsets (Figure 5A, Supplemental Figure 8B). Compared to HCs, MAS individuals had significantly more proliferating T cells and trends towards more proliferating NK cells and fewer CD56^dim^ NK cells, effector/memory B cells, and MAIT cells (Supplemental Figure 8C). However, there was considerable individual heterogeneity in MAS lymphocyte populations with a subset of MAS individuals with exceptionally large populations of effector/memory CD8^+^ T cells while one MAS individual had an increased proportion of CD56^bright^ NK cells (Supplemental Figure 8A). In addition, analysis of DEGs between MAS and HC lymphocyte subsets revealed that increased expression of the activation marker *CD38* on MAS effector/memory CD8^+^ T cells was one of the only significant DEGs between MAS and HC when corrected for multiple comparisons (Supplemental Figure 8D, Supplemental Data 4), consistent with reports that CD38 is upregulated on T and NK cells during MAS (18–20).

**Figure 5:**
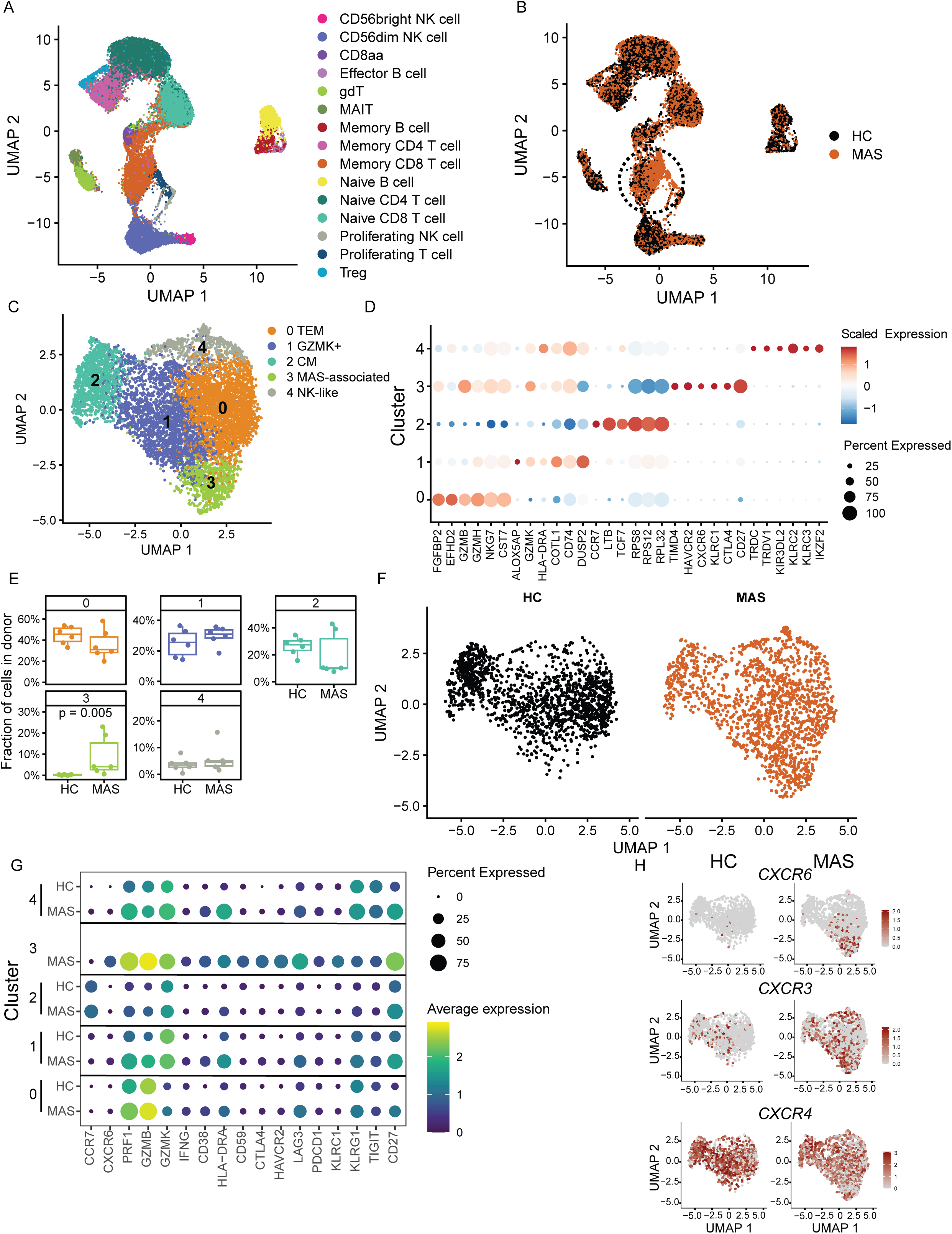
MAS-associated subset of effector/memory CD8^+^ T-cells. (A) UMAP of scRNA-Seq data from lymphocytes sorted from MAS (n=6) and HC (n=6) donors colored by cell type labels from mapping to a published dataset (27). (B) UMAP from (A) colored by study group, dotted circle shows a population of effector/memory CD8^+^ T cells populated mainly by MAS cells, downsampled to 10,000 cells per group. (C) Subclustering of effector/memory CD8^+^ T cells, colored by cluster. (D) Top six cluster-defining genes for each cluster shown as Z-score. Color indicates average scaled log-normalized expression and size of dot represents percent expressed. (E) HC and MAS cluster proportions; a Mann Whitney U test. (F) UMAPs of HC and MAS effector/memory CD8^+^ T cells, downsampled to 1,500 cells per group. (G) Expression of selected genes in scRNA-Seq data aggregated by cluster and study group. Due to the few cluster 3 cells from HCs, these are not shown. Color indicates average expression and dot size indicates the percentage of cells expressing the gene. (H) *CXCR6* (top), *CXCR3* (middle) and *CXCR4* (bottom) expression on HC (left) and MAS (right) effector/memory CD8^+^ T cells, downsampled to 1,500 cells.

Looking across all lymphocyte populations, we found *CD38* was increased primarily on NK cells and effector/memory CD8^+^ T cells (Supplemental Figure 8D). Although *HLA-DRA* appeared to be expressed on similar cells as *CD38* during MAS, the difference in *HLA-DRA* expression between MAS and HC cells was not statistically significant when corrected for multiple comparisons (Supplemental Figure 8D). As expected, we found that effector/memory CD8^+^ T cells and NK cells expressed *IFNG* transcripts during MAS (Supplemental Figure 8D), though the overall proportion of *IFNG* expressing cells was not significantly higher in CD8^+^ T cells and NK cells in MAS compared to HC.

When we compared the distribution of MAS to HC cells on the lymphocyte UMAP, we noted a region of effector/memory CD8^+^ T cells derived mainly from MAS samples (Figure 5B), suggesting specific changes to effector/memory CD8^+^ T cell compartment during MAS. To investigate this population in more detail, we subclustered effector/memory CD8^+^ T cells, which resulted in five distinct clusters (Figure 5C, Supplemental Data 5). Analysis of gene expression defined Cluster 0 as T effector memory (TEM), Cluster 1 as GZMK^+^ T cells, Cluster 2 as central memory T cells, Cluster 3 as a unique CD8^+^ T cell population in MAS, and Cluster 4 as NK-like T cells (Figure 5C-E). Of note, Cluster 3 primarily consisted of cells from MAS participants whereas the remaining four clusters included both HC and MAS cells (Figure 5E-F). The fraction of cells in Cluster 3 from MAS participants ranged from 1-23% of the effector/memory CD8^+^ T cells whereas the fraction from HCs was < 0.5% of effector/memory CD8^+^ T cells (Figure 5E); therefore, we will refer to these cells as MAS-associated CD8^+^ T cells. We also subclustered NK cells but did not find any expanded MAS-associated subcluster within the NK cells (data not shown).

We focused on the MAS-associated CD8^+^ T cells in Cluster 3 because they were uniquely expanded in MAS individuals (Figure 5E). Cluster 3 defining genes included *TIMD4, HAVCR2, CXCR6, KLRC1, CTLA4,* and *CD27* (Figure 5D). *TIMD4, HAVCR2*, *CTLA4*, and *KLRC1* encode the inhibitory receptors TIM-4, TIM-3, CTLA4, and NKG2A, which are highly expressed by chronically activated T cells found in tumors or chronic viral infections. In contrast, *CD27* is a TNF receptor family memory that provides survival signals and is usually expressed by naïve and central memory T cells, not chronically activated cells. MAS-associated CD8^+^ T cells (Cluster 3) had the highest expression of the cytolytic genes *GZMB* and *PRF1* and the highest proportion of *IFNG*-expressing cells amongst the CD8^+^ T effector/memory clusters (Figure 5G, Supplemental Figure 9), suggesting they have high effector activity.

These MAS-associated CD8^+^ T cells also expressed *CCR1*, *CCR2* and *CCR5* (Supplemental Figure 8E), which are commonly expressed on effector CD8^+^ T cells and are associated with IFNγ-producing cytotoxic CD8^+^ T cells. Interestingly, IL15, which is elevated during MAS, can upregulate CCR5 on memory CD8^+^ T cells in an antigen-independent manner (18, 43). Cluster 3 cells also expressed *CXCR6,* which is involved in T cell migration to peripheral tissues and is associated with tissue resident memory CD8^+^ T cells, and *CX3CR1*, which binds to CX3CL1 expressed by vascular endothelial cells (Figure 5G-H, Supplemental Figure 8E). The expression of both *CXCR6* and *CX3CR1* suggests that these cells may be interacting with the endothelium and migrating into tissues. Taken together, the unique appearance of Cluster 3 during MAS and the gene expression profile of these MAS-associated effector/memory CD8^+^ T cells suggest they may be active effectors poised to enter tissues during MAS.

Next, we compared the expression of select genes of interest important in CD8^+^ T cell function within CD8^+^ effector/memory subclusters between HC and MAS participants (Figure 5G and Supplemental Figure 9). Because there were so few Cluster 3 cells in HCs, we excluded them from this analysis (Figure 5G). *CD38* and *HLA-DRA* were increased in all effector/memory CD8^+^ T cells clusters during MAS (Figure 5G and Supplemental Figure 10). *CD27*, which was very highly expressed in Cluster 3 MAS-associated CD8^+^ T cells, was broadly higher in all MAS clusters, suggesting higher proliferative capacity in effector/memory CD8^+^ T cells during MAS. *PRF1* and *GZMB* were also generally higher in MAS cells than HC cells across clusters. *CXCR3*, which is expressed on activated CD8^+^ T cells and is involved in trafficking to sites of IFN-induced inflammation, was also expressed more highly across MAS CD8^+^ effector/memory clusters (Figure 5H, Supplemental Figure 8E). In contrast, *CXCR4,* which promotes CD8^+^ T cell migration to the bone marrow, was expressed similarly across all clusters with higher expression in HC as compared to MAS cells (Figure 5H, Supplemental Figure 8E). Therefore, the expression of phenotypic, trafficking, and effector molecules is broadly altered across CD8^+^ T effector/memory cells during MAS. In addition, MAS-associated CD8^+^ T cells specifically show signatures reflective of chronic activation.

Because MAS-associated CD8^+^ T cells expressed genes abundant in chronically activated T cells, we wondered whether these cells were more likely to have evidence of large monoclonal or oligoclonal T cell populations indicative of antigen-driven expansion. To investigate this question, T cell receptor (TCR) sequences were analyzed in our single-cell gene expression libraries (44). We were able to detect TCRβ CDR3 sequences in 15% of HC and 36% of MAS effector/memory CD8^+^ cells. Recovery of TCRα CDR3 was much lower so we focused on TCRβ sequences for subsequent analyses. Most of the detected TCR Vβ CDR3s were only present in a single cell (singletons; Supplemental Figure 10A). Some expanded clones were noted in Clusters 0 (TEM) and 3 (MAS-associated CD8^+^ T cells). Of the expanded clones in TEM and MAS-associated CD8^+^ T cell clusters (Clusters 0 and 3), we noted more cells represented in small (2–5) and medium (6–20) clones; there were few large clones (>20 cells with the same TCR Vβ CDR3) in any effector/memory CD8^+^ T cell cluster. When we compared MAS to HC CD8^+^ T cells, there was a trend towards more CD8^+^ TCRβ clonal expansion in MAS cells (Supplemental Figure 10B-C). Overall, while TEM and MAS-associated CD8^+^ T cell clusters showed some expanded clones, these cells were not dominated by large restricted clonal populations. Instead, effector/memory CD8^+^ T cells, including MAS-associated CD8^+^ T cells, showed clonal diversity, suggesting that non-antigen specific factors may be driving MAS CD8^+^ T cell responses.

### Cell-cell interactions during MAS

We were interested in understanding how MAS affects communication between monocytes and their interactions with lymphocytes. We used LIANA cell-cell communication analysis (45) to computationally identify important interactions between monocytes during MAS. We identified an increased enrichment in MAS monocytes in predicted interactions of *C1Q* complement components *C1QA* and *C1QB* produced by nonclassical monocytes with several C1Q receptors, including *LRP1*, expressed on all monocyte clusters (Figure 6A).

**Figure 6:**
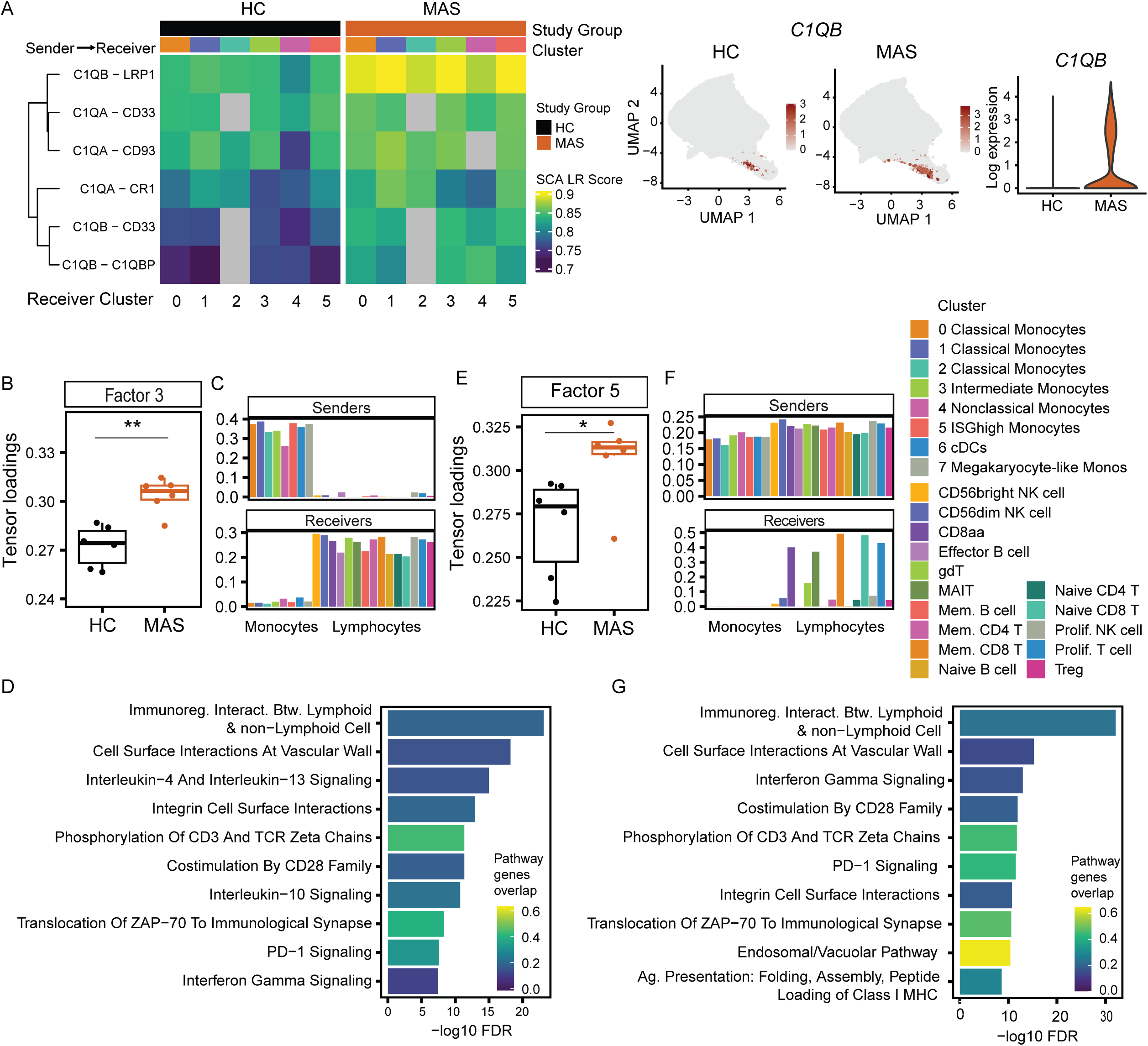
Cell-cell communication identifies altered monocyte-lymphocyte interactions in MAS. (A) On left, cell-cell communication heatmap colored by Ligand-Receptor (LR) scores for complement-associated interactions (rows) with nonclassical (cluster 4) sender cells and receiver cells grouped by monocyte cluster (columns). In middle, *C1QB* expression in HC and MAS monocytes. On right, violin plot of *C1QB* reads in nonclassical monocytes (cluster 4) from HC and MAS samples. (B-D) Tensor-cell2cell data for Factor 3 predicting monocyte to lymphocyte interactions. (B) Loading for each individual HC and MAS sample for Factor 3. (C) Loadings for sender cells (top) and receiver cells (bottom) using clusters from Figure 2B (monocytes) and Figure 5A (lymphocytes). Vertical bar plots report loadings for sender (top) and receiver (bottom) cell types and are colored by monocyte cluster or lymphocyte cell type. (D) Top enriched Reactome terms for ligands and receptors. (E-G) Tensor-cell2cell data for Factor 5 predicting all cell clusters to CD8^+^ T cell cluster interactions. (E) Loadings for each individual HC and MAS sample. (F) Loadings for sender cells (top) and receiver cells (bottom) using clusters from Figure 2B (monocytes) and Figure 5A (lymphocytes). (G) Top enriched Reactome terms for ligands and receptors.

Upregulation of *C1QB* in Cluster 4 nonclassical monocytes was consistent with this finding (Figure 6A). During MAS, there was particularly strong enrichment for *C1QB* interactions with *LRP1*, which encodes a member of the LDL receptor family that is important in scavenging material from apoptotic cells. Because C1Q proteins initiate the classical complement cascade and are opsonins for phagocytosis of apoptotic cell, these predicted interactions suggest that complement-mediated phagocytosis, perhaps of apoptotic cells, is upregulated during MAS.

We used Tensor-cell2cell to define interactions between monocyte and lymphocyte populations and determine if these interactions were significantly different between HC and MAS (46). Tensor-cell2cell reduces complex cellular communication data into key factors that capture shared patterns of signaling interactions across cell types and conditions. Six factors were identified using this analysis (Supplemental Figure 11) with factors 3 and 5 significantly higher in MAS participants compared to HCs, indicative of stronger predicted cell-cell interactions during MAS. Factor 3 consisted of monocytes as sender cells to all lymphocytes as receiver cells (Figure 6C, Supplemental Figure 11). When we evaluated the ligand-receptor pairs that defined factor 3 for enriched pathways using GSEA, we found many signaling pathways important in T cell activation and regulation including TCR signaling and CD28 co-stimulation, immunoregulatory interactions including PD-1, IL-10, and IFNγ cytokine signaling, as well as adhesion interactions such as integrins (Figure 6D). Factor 5 consisted of all clusters of leukocytes as sender cells and T cells as receiver cells (Figure 6F, Supplemental Figure 11).

Enriched pathways from the ligand-receptor pairs in factor 5 were similar to those defining monocyte-T cell interactions in factor 3 such as immunoregulatory interactions, PD-1 signaling, IFNγ signaling, and costimulation, with the addition of class I MHC antigen presentation and absence of IL-10 and IL-4/13 signaling, highlighting many of the immune functions of T cells in interacting with antigen-presenting cells. Overall, the cell-cell communication networks that were significantly higher in MAS suggest a complex picture of lymphocyte interactions and attempts at regulation.

## Discussion

In this study, we used bulk and single-cell transcriptomics to understand how monocytes and lymphocytes change during MAS. From bulk RNA-Seq, we identified a MAS monocyte gene signature score that was enriched across independent datasets and other systemic inflammatory diseases and correlated with ferritin, a known clinical biomarker of MAS. We also highlight a role for the type I IFNs, in particular IFNβ, during MAS. Although IFNγ is a well-established mediator of MAS, our findings show a significant contribution of type I IFN in influencing the transcriptional signature of all monocytes in the peripheral circulation during MAS, supported by the presence of IFNβ in circulation in active disease. In contrast to type I IFNs, we found that IFNγ principally had effects on nonclassical monocytes during MAS. Additionally, we confirmed prior observations of CD8^+^ T cell expansion during MAS and identified a population of CD8^+^ effector/memory T cells unique to MAS. Together, our findings shed light onto disease mechanisms that could guide future treatments.

In addition to showing that our MAS signature score strongly correlated with ferritin in children with sJIA and MAS in our data and a published dataset (12), we also applied our MAS monocyte signature to other systemic inflammatory diseases, cSLE and COVID-19 in adults, supporting the score’s generalizability. The correlation between the MAS signature score and SLEDAI indicated that transcriptional changes observed in monocytes from those with high disease activity in cSLE and sJIA-MAS may be driven by similar factors. This MAS signature was partly, but not entirely, driven by IFNα/β signaling; with less than 25% of the genes being known ISGs. Because the majority of the MAS signature were non-IFN related genes, our findings suggest that monocytes from active cSLE and sJIA-MAS may share other transcriptional pathways. Although MAS may develop during SLE, the dataset we used was not annotated to indicate if any individuals had SLE-MAS. Our MAS monocyte signature also included many downregulated genes, including those related to translation, which could be due to effects of type I IFN signaling as part of antiviral defense.

We found that the dominant signature during MAS across all classical monocyte clusters was from IFNα/β rather than IFNγ and identified a population of ISG^high^ monocytes expanded during MAS and driven by IFNα/β signaling. This was surprising given the importance placed on the role of IFNγ during MAS and other forms of HLH. Our findings suggest a potentially important and previously underrecognized role for type I IFN during MAS. While IFNγ contributed to the transcriptional signature of nonclassical monocytes during MAS in our study, it played little role in shaping the transcriptional signature of the ISG^high^ classical monocyte population. Importantly, we found elevated circulating IFNβ during MAS. Few prior studies have suggested a role for IFNα/β signaling during MAS (18, 47). Although Huang and colleagues also found evidence for type I IFN activity during MAS using an IFN reporter assay (18), here we showed direct detection of IFNβ from MAS plasma using an ultrasensitive IFN detection assay. Notably, levels of IFNβ during MAS were higher than in active sJIA without MAS. We note that the highest IFNβ detected from the quiescent group was from a participant with sJIA-associated lung disease, which suggests ongoing subclinical inflammation that may contribute to lung pathology. Similarly, Marques et al. found an elevated type I IFN signature was associated with sJIA-lung disease (48). Taken together, these data suggest that IFNβ drives transcriptional changes in MAS monocytes and may have an underappreciated role in the etiopathogenesis of MAS.

What is the source of IFNβ during MAS? Many hematopoietic and non-hematopoietic cells can produce IFNβ, including macrophages, DCs, fibroblasts, and endothelial cells. We did not detect transcripts encoding IFNβ or any of the IFNαs in our bulk or single-cell RNA-Seq data. However, given the low sensitivity of scRNA-Seq and the low amounts of these cytokines in circulation, it is still possible that monocytes produce IFNα/β during MAS that was below the limits of detection in our assays. Alternatively, other cells in tissue or the vasculature may produce these cytokines. Because we did not find circulating IFNα2 in the majority MAS participants, we also presume that pDCs are not the principal source of IFNα/β, however we do not exclude the possibility that in the few individuals with circulating IFNα2 that IFNs may come from pDCs. Whether other myeloid cells, such as cDCs, tissue macrophages, or neutrophils produce IFNβ during MAS, or whether non-hematopoietic cells are the main source, is an important future question.

What signals induce IFNβ production during MAS is also an open question. IFNβ is principally produced in response to innate sensing of viral nucleic acids during infection or self-nucleic acids from dead or dying cells, and many ISGs, including some found in our MAS signature score or upregulated in our scRNA-Seq data, encode viral sensors, such as *ZNFX1*, or molecules that regulate viral sensors, such as *TRIM25*. Given that primary HLH and other forms of sHLH are often induced by viral infection, these findings suggest that MAS during sJIA may also be triggered by viruses (6, 35, 49). Alternatively, self-nucleic acids recognized after phagocytosis of apoptotic cells may induce IFNβ during MAS. This is an intriguing possibility as we saw strong upregulation of *C1QB*, an opsonin for phagocytic cells, and predicted C1Q interactions between monocytes during MAS in our scRNA-Seq analyses. Similarly, *C1QA-C* transcripts are induced in monocytes from patients with sJIA lung disease (17). Future work will aim to understand the innate sensors, the sources of their DNA/RNA ligands, and the role of C1Q during MAS.

We and others have identified expansion of proliferating T cells in individuals with MAS (18). In addition, using our scRNA-Seq analyses, we defined a unique MAS-associated CD8^+^ T cell effector/memory subpopulation that was specifically associated with MAS. These MAS-associated CD8^+^ T cells had a signature of cytotoxic potential with high expression of perforin and granzyme B, similar to that of CD8^+^ TEM cells, along with expression of *CCR1*, *CCR2* and *CCR5*. However, these cells also expressed transcripts encoding multiple inhibitory receptors associated with chronically activated T cells seen in viral infections and in tumors, including PD1, CTLA4, LAG3, and TIM-3, and expressed *CD27,* which is not typically expressed on chronically activated or exhausted T cells but rather is typically associated with proliferative and effector-like T cells. Interestingly, a similar phenotype has been described in islet autoantigen specific CD8^+^ T cells in type 1 diabetes (50), suggesting that these cells may be present in multiple autoinflammatory or autoimmune settings. MAS-associated CD8^+^ T cells also expressed elevated levels of *CXCR6*, associated with tissue T cells, and *CX3CR1*, which allows for interactions with vascular endothelial cells, suggesting that these CD8^+^ T cells are destined to leave the bloodstream and enter tissues. This is supported by our Tensor-cell2cell analysis that also enriched for association of T cells with vascular endothelium during MAS. Interestingly, these IFNγ-producing CD8^+^ T cells and nonclassical monocytes that exhibited an IFNγ signature both expressed *CX3CR1* perhaps suggesting physical proximity between these cells at the endothelium may account for why nonclassical monocytes demonstrated an IFNγ signature while other monocytes did not. Preliminary TCR analyses suggested that this population was not clonally expanded as might be expected from an antigen-driven response. Therefore, these cells may instead be driven by the inflammatory cytokine milieu or cell-cell interactions. Detailed study of these cells warrants further investigation in the future.

Interestingly, in our cell-cell communication analysis, we noted that immunoregulatory interactions between monocytes and lymphocytes dominated the cell-cell interactions, suggesting that circulating immune cells attempt to reduce hyperactivity during MAS. Similar to others (51, 52), we found increased concentrations of IL-10 in MAS plasma despite high inflammation. This raises the question of whether MAS cells are insensitive to IL-10 signaling and other immunoregulatory strategies or, alternatively, whether immunoregulatory interactions may instead have proinflammatory effects in the MAS environment. For example, when cells are primed with type I IFN, cellular responses to subsequent IL-10 signaling can include enhanced STAT1 activation (53). This potential pro-inflammatory role of IL-10 during MAS is further supported by mouse model data in which IL-10 blockade protected animals from lethal HLH (51), however other MAS models require IL-10 blockade for full disease (54). This raises the intriguing question of how the inflammatory milieu may modulate immune signaling during MAS.

The limitations of this study include the small number of participants, heterogeneity in treatments and treatment duration, and the small number of completely untreated participants. On the other hand, a strength of our study was analysis of samples from participants prior to corticosteroid treatment which we showed had substantial effect on gene and protein expression. Despite these limitations, we did identify intriguing findings that warrant further investigation, including a strong role for type I IFN and unique CD8^+^ T cell phenotypes during MAS. Future studies with larger numbers of untreated participants and further longitudinal tracking of response to therapies in individuals will be informative. In addition, our study did not include TCR sequencing to formally assess for T cell clonality during MAS. Our limited analyses of our scRNA-Seq data for TCR Vβ segments did not suggest clonal expansion, and several other studies have similarly suggested that T cells in MAS are likely not clonally expanded (18, 20). However, future analysis of TCR using TCR-Seq would be the most definitive way to evaluate for TCR clonality.

In conclusion, we demonstrated that MAS monocytes had specific transcriptional changes, many of which were driven by IFNβ. These included expansion of an ISG^high^ monocyte population, upregulation of CD16 on classical monocytes, and broad transcriptional changes observed across the monocyte and lymphocyte compartments. Notably, we found circulating IFNβ in the plasma of MAS participants prior to initiation of systemic corticosteroid treatment, highlighting its potential role early in disease pathogenesis. In addition, we identified altered monocyte-monocyte and monocyte-lymphocyte interactions during MAS, involving molecules associated with antigen presentation, cell trafficking, and immunoregulation. These findings provide new insights into monocyte-driven mechanisms during MAS and suggest novel avenues for therapeutic targeting, including the potential use of therapies targeting type I IFNs or IFNα/β signaling.

## Methods

### Study groups

Children with suspected or confirmed sJIA or MAS were recruited at Seattle Children’s Hospital or Cohen Children’s Medical Center, Northwell Health and all diagnoses were defined based on the treating clinician’s assessment. MAS secondary to sJIA was confirmed using the 2016 ACR/EULAR MAS Classification Criteria for MAS complicating sJIA (5) and a ferritin/ESR ratio of greater than 21.5 (55). Quiescent disease were patients without fever, rash, arthritis, or notable laboratory abnormalities. HC samples were from the Benaroya Research Institute’s Healthy Control Biorepository. Study data were collected using REDCap electronic data capture tools hosted at the University of Washington and Northwell Health.

Demographics and clinical and laboratory data of participants are provided in Supplemental Tables 2-5. Ancestry and ethnicity data were self-identified by the participants. Clinical and laboratory data were extracted from medical records.

### Sex as a biological variable

Both male and female participants were included, and HCs were sex matched to study participants.

### Sample collection

Individuals with known or suspected sJIA with or without features of MAS were consented for blood collection in coordination with clinical lab draws. Blood was collected and transported to Benaroya Research Institute or the Feinstein Institutes for Medical Research for processing. PBMCs and plasma were isolated using Ficoll-Paque PLUS (Cytiva) according to the manufacturer’s instructions. Cells were frozen and stored in liquid nitrogen and plasma stored at -80C until analyzed.

### Flow cytometry and cell sorting

Antibodies and sources are listed in Supplemental Table 6. Thawed PBMCs were stained with the live/dead-APC-E780 (eBiosciences) for 30 minutes at 4C, followed by antibody staining for 30 minutes at 4C. For bulk RNA sequencing, CD14^+^ monocytes were isolated by gating on live, CD3^-^CD19^-^CD56^-^CD15^-^CD14^+^HLA-DR^+^ cells (Supplemental Figure 2A) on a FACSAria Fusion (BD Biosciences). The CD14 gate was set based on a sample stained for all antibodies except CD14-BV510. For scRNA-Seq, monocytes were isolated by gating on live, CD3^-^CD19^-^CD56^-^CD14^+^ cells (Supplemental Figure 5) on a FACSAria Fusion (BD Biosciences). Lymphocytes were isolated by gating on live, CD3^+^CD19^+^CD56^+^CD14^-^ cells (Supplemental Figure 5) on a FACSAria Fusion (BD Biosciences). All cells were collected in media containing 10% human AB serum (Omega Scientific). Prior to sorting, cells were labeled with hashtags (Biolegend TotalSeq-C).

### Generation of IFNβ and IFNγ stimulated monocytes

CD14^+^ monocytes were isolated from PBMCs from five females (ages 14-30) using CD14 microbeads (Miltenyi) and cultured in RPMI-1640 (Cytiva) with 10% human AB serum (Omega Scientific). Cells were treated with 100 U/ml IFNβ (PBL) or 25 ng/ml IFNγ (Peprotech) or in media alone for 24 hours. RNA was isolated using RNeasy Micro kit (Qiagen) and analyzed by bulk RNA-Seq.

### Bulk RNA-Seq of CD14^+^ monocytes

RNA from sorted monocytes was isolated using RNeasy Micro kit (Qiagen). Libraries were prepared using a SMART-Seq v4 Ultra Low Input RNA Kit for Sequencing (Takara Bio USA, Cat #634891) and Nextera XT DNA Library Preparation Kit (Illumina, FC-131-1096). Libraries were pooled, quantified by a Qubit Fluorometer (Life Technologies) and sequenced on a NextSeq 2000 sequencer (Illumina) with paired-end 59bp targeting 5 million reads/sample. FASTQ files were generated in BaseSpace (Illumina) and processed on the Galaxy platform (56). Reads were trimmed (FASTQ Trimmer tool v1.0.0), adapters filter (FastqMcf v1.1.2) and aligned to GRCh38 with gene annotations from Ensembl release 91 (STAR aligner v2.4.2a) (57, 58). Gene counts were generated using HTSeq-count (v0.4.1) (59). Quality metrics were compiled using Picard (v1.134), FastQC (v0.11.3), Samtools (v1.2) (60), and HTSeq-count. Samples with > 5 × 10^5 FASTQ reads, > 55% mapped reads, and median CV coverage < 1.4 were retained for analysis.

### Differential expression and MAS signature score generation

Differential expression analysis was performed in R using limma package with the voomWithQualityWeights transformation of TMM-normalized counts (v3.6.62) (61). A MAS signature score was calculated as the average log2 expression of genes upregulated in MAS minus the average expression of downregulated genes (FDR < 0.05).

### Analysis of monocytes from adults hospitalized with COVID-19

Classical monocytes were isolated from frozen PBMCs (HC n=10, adults hospitalized with COVID n=26) using cell sorting and analyzed using RNA-Seq as described in (25). A t-test was used to compare the MAS signature score between the two groups.

### Analysis of public RNA-Seq data

Bulk RNA-Seq counts from the NCBI GEO dataset GSE147608 containing pediatric controls, sJIA cases, and MAS cases were analyzed (12). scRNA-Seq profiles from pediatric SLE cases and controls from the NIH dbGaP study phs002048.v2.p1 were also analyzed (24). MAS signature scores within monocytes were compared with ferritin or SLEDAI scores, respectively.

#### scRNA-Seq of monocytes and lymphocytes

Sorted monocytes and lymphocytes were sequenced across three batches using 10x Chromatin X with target captures of 10,000-20,000 cells/channel. Libraries were prepared using Chromium Next GEM Single Cell 3’ v1, 5’ v2, or 5’ GEM-X kits and sequenced on a NextSeq 2000 (Illumina) with a target capture of 10,000 for monocytes (batches 1-2), 13,000 for lymphocytes (batch 2) or 20,000 for monocytes and lymphocytes (batch 3) per channel. Base calls were processed to FASTQ files on BaseSpace.

Demultiplexed reads were processed using Cell Ranger (v7.1.0, 10x Genomics) and analyzed in Seurat (v5.0.0) (62). Standard preprocessing included normalization, variable gene selection, scaling, PCA, Louvain clustering, and UMAP visualization (63). Cluster markers were identified with FindAllMarkers, Hashtag-based donor assignment was performed manually, and cell types were annotated using CellTypist (64) with signatures derived from human PBMCs (27).

### Pseudobulk analysis of scRNA-Seq data

Pseudobulked differential expression analysis was performed on cells aggregated by donor and cluster and modeling disease group and batch effects using DESeq2. Wald tests with FDR correction were used for significance testing (v1.38.3) (65).

### Pathway analysis

Pathway enrichment analysis was performed using the enrichR package (v3.2) (66) with the Reactome 2022 database. Interferon signatures were evaluated using monocyte-specific IFNβ or IFNγ response signatures and non-negative least squares factorization (41). Within clusters, AUCell (v3.21) was used to compare interferon signatures of MAS and HC groups (67).

### Measurement of human cytokines in plasma

The Human ProInflammatory-9 S-Plex Ultra-Sensitive Kit (K15396S), IFN-α2α S-Plex Ultra Sensitive Kit (K151P3S) and IFN-β S-Plex Ultra Sensitive Kit (K151ADRS) from MSD were used to measure plasma IL-1β, IL-6, IL-10, IL-12p70, TNF, IFNγ, IFNα2a and IFNβ. All three assays were performed simultaneously according to the manufacturers’ instructions and utilized a Tecan Fluent 1080 for all pipetting steps to allow for concurrent processing. Samples were thawed in a 37°C water bath, and centrifuged for 3 minutes at 2000g. For samples with less than 125 μL volume, samples were diluted in MSD kit supplied diluent 58 to a final volume of 125 μL and 25 μL was used per assay plate. MSD plates were analyzed on the Meso Sector S 600MM (MSD) and data processed with MSD Discovery Workbench Software. Lower limit of detection (LLOD) was defined as the concentration of diluted standard that provides signal 2.5 standard deviations above the mean of the blank value. Lower limit of quantification (LLOQ) was defined as the lowest concentration of diluted standard with a coefficient of variation (CV) less than 20% between replicates.

### TCR analysis

TCR sequences were inferred from single-cell libraries using TRUST4 (44). Productive TCR α and β chain sequences were identified from the gene expression data. We recovered TCRα CDR3 sequences in 12% and TCRβ sequences in 31% of effector/memory CD8^+^ T cells.

### Cell-cell interaction analyses

Cell-cell interaction analysis was performed using Liana (v0.1.14) (68) and Tensor-cell2cell (v0.7.4) (46). Aggregated statistics incorporated NATMI, Connectome, iTALK, and SingleCellSignalR. Tensor-cell2cell identified an optimal rank of six. Pathway enrichment was performed on ligand and receptor loadings of factors larger than a cutoff of 0.005 using enrichR with Reactome 2022.

### Study Approval

The study was approved by the Institutional Review Boards of Seattle Children’s Hospital, Benaroya Research Institute, and Northwell Health.

**Table 1:**
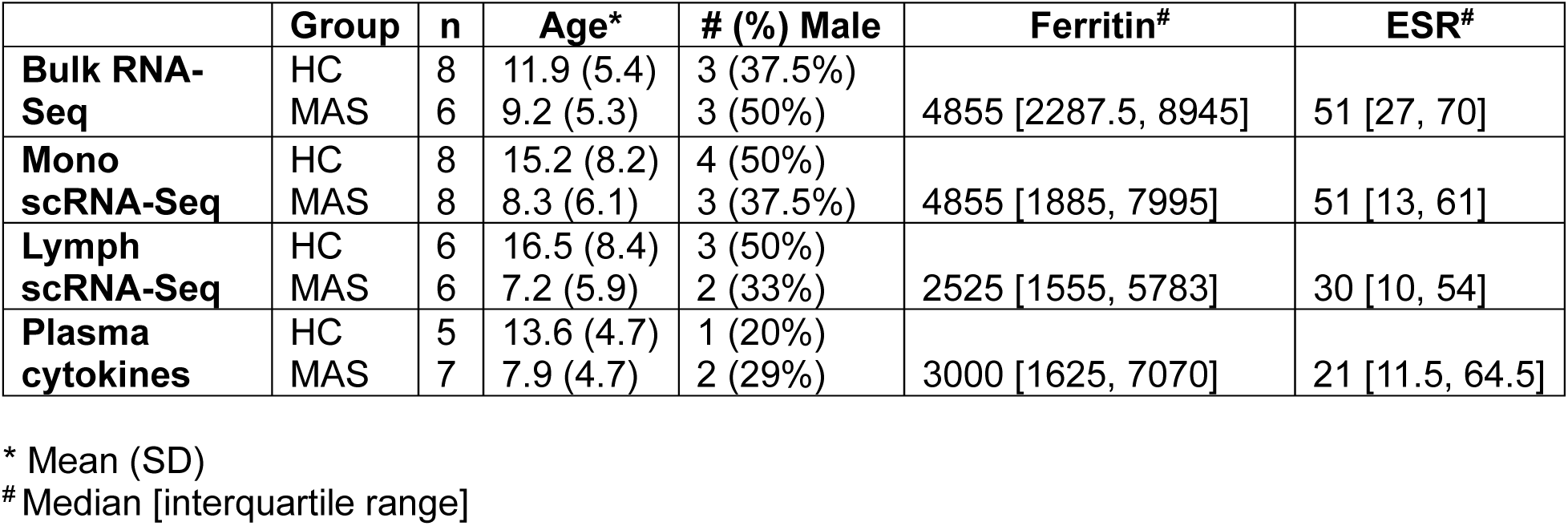
Summary of participant demographics.

## Data Availability

Sequencing data from MAS cohort be submitted to dbGAP. Bulk RNA-Seq of CD14^+^ monocytes from COVID-19 cohort are deposited in GEO (GSE334777).

## Author Contributions

Conceptualization: SPC, JAH, HAD, ELK, SS, JHY, BJB, DJC, CM, CS

Data curation: HAD, NM

Formal analysis: SPC, HAD, ELK, NM

Funding acquisition: JAH, BJB, SPC

Investigation: SPC, HAD, ELK, NM, GG, AL, AH, ARO, EDL, PB, PJW

Resources: SS, JHY, CS

Writing – original draft: SPC, HAD, JAH, ELK, DJC, BJB

Writing – review & editing: all authors

## Acknowledgements

The authors thank the study participants and their families for their participation and the clinicians at Seattle Children’s Hospital and Cohen Children’s Medical Center caring for these patients. The authors thank the members of the Division of Rheumatology in the Department of Pediatrics at the University of Washington and Division of Pediatric Rheumatology in the Department of Pediatrics at Cohen Children’s Medical Center, Northwell Health. We thank Dr. Yongdong Zhao for management of Seattle Children’s rheumatology biorepository. The authors appreciate the assistance of clinical research coordinators, including Mary Eckert, Michelle Geiszler, Avery Hubert, Ching Hung, Andrea Connolly, Nidhi Naik, and coordinators and staff in the BRI Center for Interventional Immunology for patient recruitment and data collection. The authors gratefully acknowledge the contributions of Catalina Sakai, Mackenzie S. Kopp, Saransh N. Kaul, and Troy R. Torgerson of the Allen Institute, Immunology for collaboration on the analyses presented. The authors gratefully acknowledge the expertise of the Benaroya Research Institute’s Clinical Core Laboratory, including Thien-Son Nguyen, Adam Wojno of the Cell and Tissue Analysis Group, Vivian Gersuk, Quynh-Anh Nguyen, and Kimberly O’Brien of the Genomics Core, and members of the Hamerman lab, especially Susana Orozco, Natalie Thulin, and Hayley Waterman, and members of the Barnes lab for helpful discussions. The authors thank Dr. Anne Hocking for reviewing and editing the manuscript.

