## Supplemental Tables and Figures for "Type I Interferon-Driven Monocyte Dysregulation and MAS-associated CD8^+^ T cells During Macrophage Activation Syndrome"

Supplemental Table 1: Participant Information

| Sample Number | Group | Bulk RNA-Seq | Monocyte scRNA-Seq | Lymphocyte scRNA-Seq | Cytokine Quant |
|---|---|---|---|---|---|
| sJIA1 | Inactive<br>MAS | + | + | + | +<br>+ |
| sJIA2 | MAS<br>Inactive | +<br>+ | + | + | +<br>+ |
| sJIA3 | sJIA<br>Inactive<br>Inactive | +<br>+ | | | +<br>+<br>+ |
| sJIA4 | Inactive | + | | | + |
| sJIA5 | Inactive | + | | | + |
| sJIA6 | Inactive | + | | | + |
| sJIA7 | MAS | | + | + | + |
| sJIA8 | sJIA<br>Inactive | +<br>+ | | | +<br>+ |
| sJIA9 | MAS | + | | | + |
| sJIA10 | MAS<br>Inactive | +<br>+ | + | + | +<br>+ |
| sJIA11 | MAS<br>Post IVMP* | +<br>+ | + | | |
| sJIA12 | MAS | + | + | + | + |
| sJIA13 | sJIA | + | | | + |
| sJIA14 | MAS<br>Post IVMP* | +<br>+ | + | | |
| sJIA15 | sJIA | + | | | + |
| sJIA16 | MAS | | + | + | |
| sJIA17 | MAS | | | | + |
| sJIA18 | sJIA | | | | + |
| HC1 | HC | + | + | + | <br>+ |
| HC2 | HC | + | | | |
| HC3 | HC | + | | | |
| HC4 | HC | + | + | | |
| HC5 | HC | + | | | + |
| HC6 | HC | + | + | | + |
| HC7 | HC | + | | | + |
| HC8 | HC | + | | | |
| HC9 | HC | | + | + | |
| HC10 | HC | | + | + | |
| HC11 | HC | | + | + | + |
| HC12 | HC | | + | + | |
| HC13 | HC | | + | + | |

* Post IVMP = sample collected after three days of IV methylprednisolone pulses (30 mg/kg/dose to a max of 1 gram)

Supplemental Table 2: Clinical demographic and laboratory values for bulk CD14+ monocyte RNA sequencing

| Group | Healthy | Quiescent | Active sJIA | Active MAS |
|---|---|---|---|---|
| **Number** | 8 | 8 | 4 | 6 |
| **Mean age** in years (SD) | 11.9 (5.4) | 12.3 (5.3) | 9 (7.7) | 9.2 (5.3) |
| **# Males** (% of group) | 3 (37.5%) | 3 (37.5%) | 2 (50%) | 3 (50%) |
| **Race** – | | | | |
| American Indian | 0 (0%) | 0 (0%) | 0 (0%) | 0 (0%) |
| Asian | 1 (12.5%) | 2 (25%) | 1 (25%) | 1 (16.7%) |
| Black | 0 (0%) | 0 (0%) | 0 (0%) | 0 (0%) |
| White | 7 (87.5%) | 6 (75%) | 3 (75%) | 5 (83.3%) |
| Unknown | 0 (0%) | 0 (0%) | 0 (0%) | 0 (0%) |
| **Ethnicity** – Hispanic # (%) | 1 (12.5%) | 2 (25%) | 0 (0%) | 0 (0%) |
| **Laboratory Studies*** | not done | | | |
| WBC (x$10^3$/mL) | | 5.5 [5.1, 8.0] | 11.5 [9.9, 12.6] | 9.1 [9, 9.5] |
| Hgb (g/dL) | | 11.4 [10.6, 14.1] | 11.4 [10.6, 11.6] | 9.3 [9.0, 10.8] |
| Platelets (x$10^3$/mL) | | 329 [280.1, 361.8] | 427.5 [330, 493.5] | 400.5 [350.8, 454] |
| CRP (mg/dL) | | 0.1 [0.1, 0.4] | 3.4 [1.8, 5.1] | 11 [7.2, 14.6] |
| ESR (mm/hr) | | 4 [2, 22] | 78.5 [63.8, 88] | 51 [27, 70] |
| Ferritin (ng/mL) | | 44 [17.5, 57.5] | 477.5 [144.3, 835.8] | 4855 [2287.5, 8945] |
| AST (IU/L) | | 20.5 [19.8, 33.5] | 42 [38, 46.5] | 63 [62, 100] |
| Fibrinogen (mg/dL) | | 311 [260, 355] | 451 [398.5, 585.5] | 367 [354, 499] |
| LDH (IU/L) | | 260 [206.5, 333] | 590 [530.5, 630.5] | 2015 [1564.5, 2077] |
| **Duration of disease**** | n/a | 985 [672, 1442] | 429 [3, 2021] | 1 [1, 3] |
| **Medications** | | | | |
| None | | 2 | 1 | 2 |
| Steroids | | 2 | 2 | 0 |
| IL-1 inhibitor | | 4 | 1 | 4 |
| IL-6 inhibitor | | 1 | 1 | 0 |
| Methotrexate | | 1 | 0 | 1 |
| Calcineurin Inhibitor | | 1 | 0 | 0 |
| TNF inhibitor | | 1 | 1 | 0 |
| Other | | 4 | 2 | 0 |

* Laboratory studies: median and interquartile range

** Duration of disease in days (median, interquartile range) at time of blood draw from diagnosis

Supplemental Table 3: Clinical demographic and laboratory values for monocyte single cell RNA sequencing

| Group | Healthy | Active MAS |
|---|---|---|
| Number, % of total | 8 (50%) | 8 (50%) |
| **Mean age** in years (SD) | 15.1 (8.2) | 8.3 (6.1) |
| **# Males** (% of group) | 4 (50%) | 3 (37.5%) |
| **Race** – | | |
| American Indian | 0 (0%) | 0 (0%) |
| Asian | 0 (0%) | 1 (12.5%) |
| Black | 0 (0%) | 0 (0%) |
| White | 8 (100%) | 7 (87.5%) |
| Unknown | 0 (0%) | 0 (0%) |
| **Ethnicity** – Hispanic #, % | 0 (0%) | 0 (0%) |
| **Laboratory Studies*** | not done | |
| WBC (x$10^3$/mL) | | 9 [6, 11] |
| Hgb (g/dL) | | 11 [9, 12] |
| Platelets (x$10^3$/mL) | | 401 [303, 453] |
| CRP (mg/dL) | | 6 [5, 10] |
| ESR (mm/hr) | | 51 [13, 61] |
| Ferritin (ng/mL) | | 4855 [1885, 7995] |
| AST (IU/L) | | 88 [56, 135] |
| Fibrinogen (mg/dL) | | 361 [300, 417] |
| LDH (IU/L) | | 1326 [414, 1994] |
| **Duration of disease** | | |
| Median in days from diagnosis | | 9 [1, 303] |
| **Medications** | N/A | |
| None | | 3 |
| Steroids | | 0 |
| IL-1 inhibitor | | 5 |
| IL-6 inhibitor | | 0 |
| Methotrexate | | 1 |
| Calcineurin inhibitor | | 1 |
| Other | | 1 |

*Laboratory studies: median and interquartile range

Table 4: Clinical demographic and laboratory values for lymphocyte single cell RNA sequencing

| Group | Healthy | Active MAS |
|---|---|---|
| Number, % of total | 6 (50%) | 6 (50%) |
| **Mean age** in years (SD) | 16.5 (8.4) | 7.2 (5.9) |
| **# Males** (% of group) | 3 (50%) | 2 (33%) |
| **Race** – | | |
| American Indian | 0 (0%) | 0 (0%) |
| Asian | 0 (0%) | 1 (17%) |
| Black | 0 (0%) | 0 (0%) |
| White | 6 (100%) | 5 (83%) |
| Unknown | 0 (0%) | 0 (0%) |
| **Ethnicity** – Hispanic #, % | 0 (0%) | 0 (0%) |
| **Laboratory Studies*** | not done | |
| WBC (x10$^3$/mL) | | 7.7 [5.6-9.2] |
| Hgb (g/dL) | | 11.1 [9.7-11.6] |
| Platelets (x10$^3$/mL) | | 448 [233.5-455.5] |
| CRP (mg/dL) | | 5.3 [3.9-6.4] |
| ESR (mm/hr) | | 30 [10-54] |
| Ferritin (ng/mL) | | 2525 [1555-5783] |
| AST (IU/L) | | 75 [52-1450] |
| Fibrinogen (mg/dL) | | 341 [243-381] |
| LDH (IU/L) | | 818 [413-1779] |
| **Duration of disease** | | |
| Median in days from diagnosis | | 48 [5-749] |
| **Medications** | N/A | |
| None | | 2 |
| Steroids | | 0 |
| IL-1 inhibitor | | 4 |
| IL-6 inhibitor | | 0 |
| Methotrexate | | 1 |
| Calcineurin inhibitor | | 1 |
| Other | | 1 |

*Laboratory studies: median and interquartile range

Supplemental Table 5: Clinical demographic and laboratory values for cytokine analysis

| Group | Healthy | Quiescent | Active sJIA | Active MAS |
|---|---|---|---|---|
| **Number** | 5 | 8 | 5 | 7 |
| **Mean age** in years (SD) | 13.6 (4.2) | 11.6 (5.7) | 9.2 (6.6) | 7.9 (4.7) |
| **# Males** (% of group) | 1 (20%) | 3 (37.5%) | 2 (40%) | 2 (29%) |
| **Race** – | | | | |
| American Indian | 0 (0%) | 0 (0%) | 1 (20%) | 0 (0%) |
| Asian | 2 (40%) | 1 (12.5%) | 0 (0%) | 1 (14%) |
| Black | 0 (0%) | 0 (0%) | 0 (0%) | 0 (0%) |
| White | 3 (60%) | 7 (87.5%) | 4 (80%) | 5 (71%) |
| Unknown | 0 (0%) | 0 (0%) | 0 (0%) | 1 (14%) |
| **Ethnicity** – Hispanic #, % | 0, 0% | 2, 25% | 1, 20% | 0, 0% |
| **Laboratory Studies*** | not done | | | |
| WBC (x$10^3$/mL) | | 5.8 [5.2, 6.7] | 10.6 [9.3, 12.4] | 9 [5.9, 9.1] |
| Hgb (g/dL) | | 12 [10.6, 14] | 11.2 [9.8, 11.5] | 10.5 [9.3, 11.6] |
| Platelets (x$10^3$/mL) | | 328 [287, 345] | 374 [362, 493] | 451 [400, 512] |
| CRP (mg/dL) | | 0.1 [0.1, 0.4] | 4.6 [2.1, 6.6] | 6.7 [5.3, 10.1] |
| ESR (mm/hr) | | 2 [2, 20] | 77 [71, 80] | 21 [11.5, 64.5] |
| Ferritin (ng/mL) | | 48 [19, 67] | 783 [172, 994] | 3000 [1625, 7070] |
| AST (IU/L) | | 21 [20, 36] | 44 [40, 54] | 63 [55.5, 153] |
| Fibrinogen (mg/dL) | | 288 [255, 345] | 562 [425, 684] | 1965 [414, 2073] |
| LDH (IU/L) | | 290 [213, 363] | 563 [433, 684] | 372 [250, 420] |
| **Duration of disease**** In days | | 1072 [698, 1363] | 3 [1, 854] | 15 [2.5, 526] |
| **Medications** | N/A | | | |
| None | | 2 | 1 | 1 |
| Steroids | | 2 | 2 | 1 |
| IL-1 inhibitor | | 4 | 2 | 6 |
| IL-6 inhibitor | | 1 | 1 | 0 |
| Methotrexate | | 1 | 0 | 1 |
| Calcineurin inhibitor | | 1 | 0 | 1 |
| TNF inhibitor | | 1 | 1 | 0 |
| Other | | 4 | 3 | 1 |

*Laboratory studies: median and interquartile range
** Duration of disease in days (median, interquartile range) at time of blood draw from diagnosis

Supplemental Table 6: List of Flow Cytometry Antibodies and Sources

| Target Antigen | Clone | Conjugate | Source |
|---|---|---|---|
| CD14 | M5E2 | BV510 | Biolegend |
| CD16 | 3G8 | PE | Biolegend |
| CD15 | W6D3 | BV650 | Biolegend |
| CD3 | OKT3 | BV650 | Biolegend |
| CD19 | HIB19 | BV650 | Biolegend |
| CD56 | 5.1H11 | BV650 | Biolegend |
| HLA-DR | L243 | APC | Biolegend |
| CD163 | GH1/61 | PE-Cy7 | Biolegend |

Supplemental Figure 1

A. Bulk RNA-Seq of CD14⁺ Monocytes

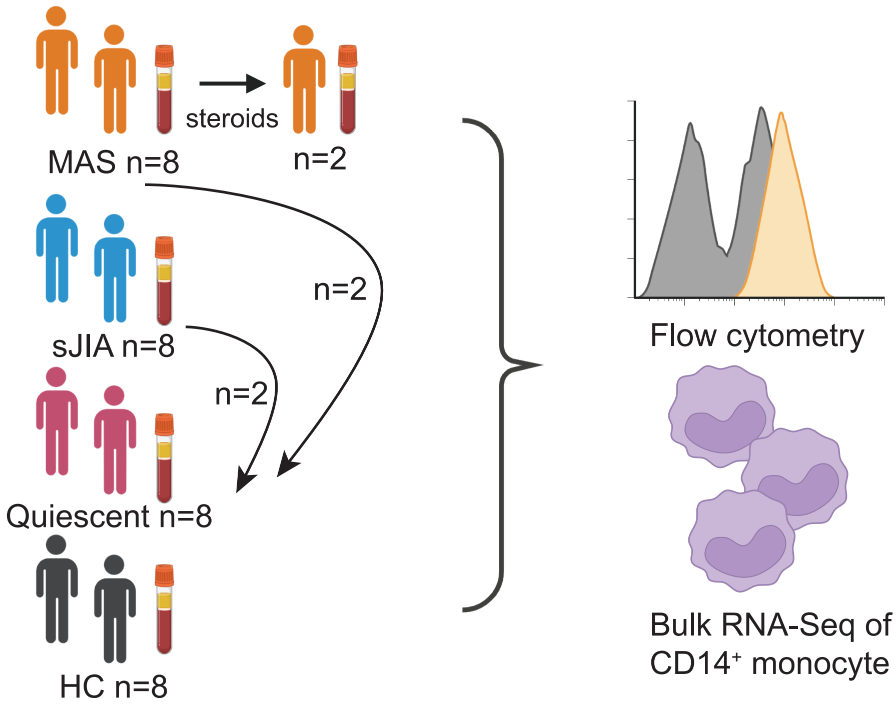


B. scRNA-Seq

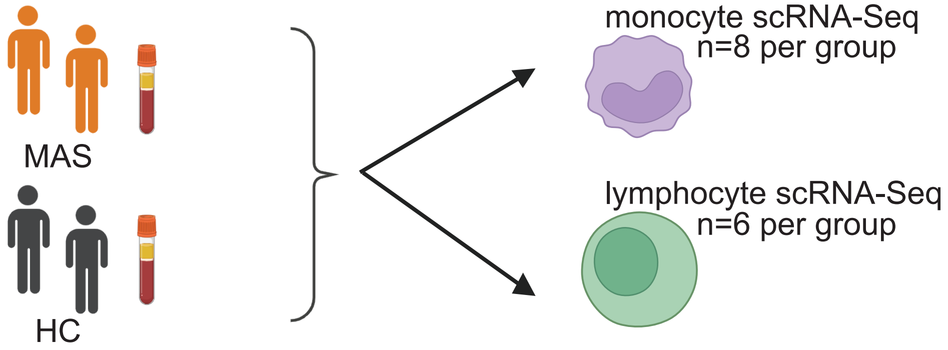


C. Plasma Cytokines

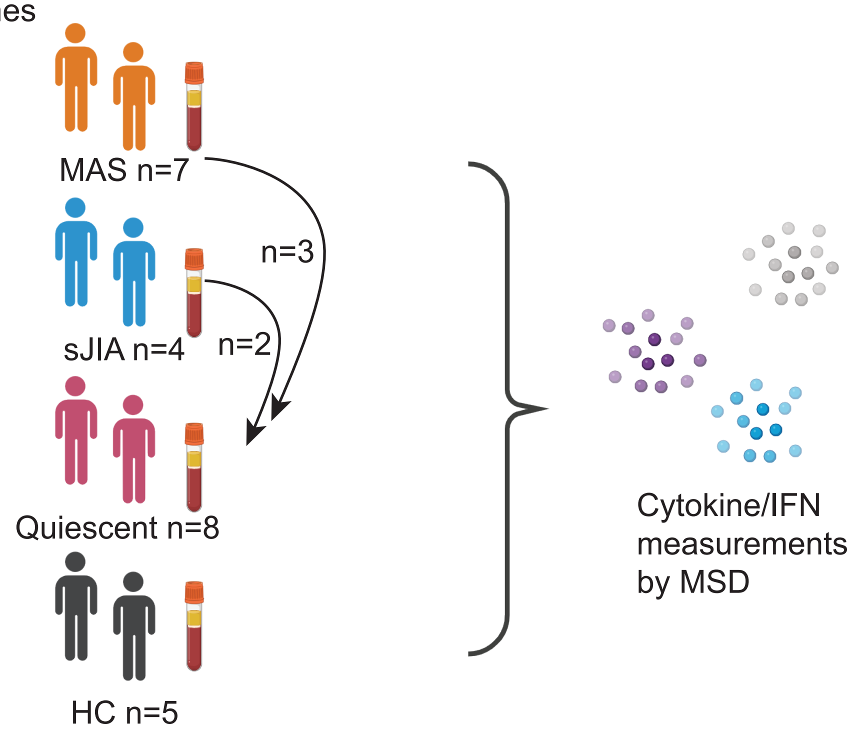

**Supplemental Figure 1: Schematic of experimental design.** (A) Groups of participants that provided PBMCs for monocyte bulk RNA-Seq of classical monocytes and flow cytometry assessment of monocytes. (B) Groups of participants that provided PBMCs for scRNA-Seq of monocytes and lymphocytes. (C) Groups of participants that provided plasma for cytokine assessment. Arrows indicate participants that provided longitudinal samples from different disease states.

Supplemental Figure 2

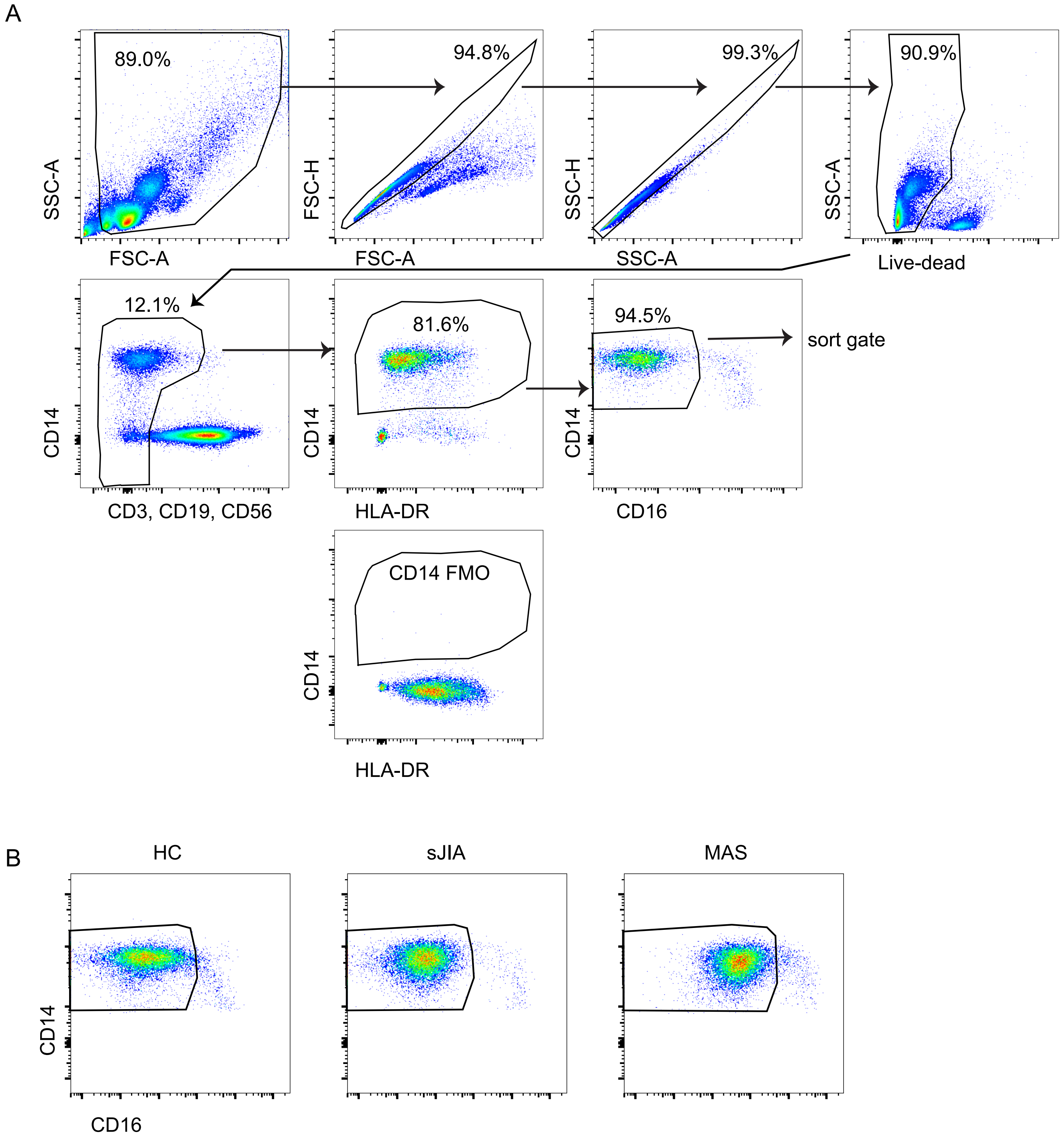

**Supplemental Figure 2: Gating for sorting monocytes for bulk RNA-Seq.** (A) PBMCs were stained with the indicated antibodies and gated as shown to identify $CD14^{+}CD16^{-/low}$ live monocytes. At bottom is FMO for CD14 which was used on a consistent control sample for each day of sorting to set $CD14^{+}$ gate. (B) Representative CD14 x CD16 plots from HC, sJIA, and MAS individuals.

Supplemental Figure 3

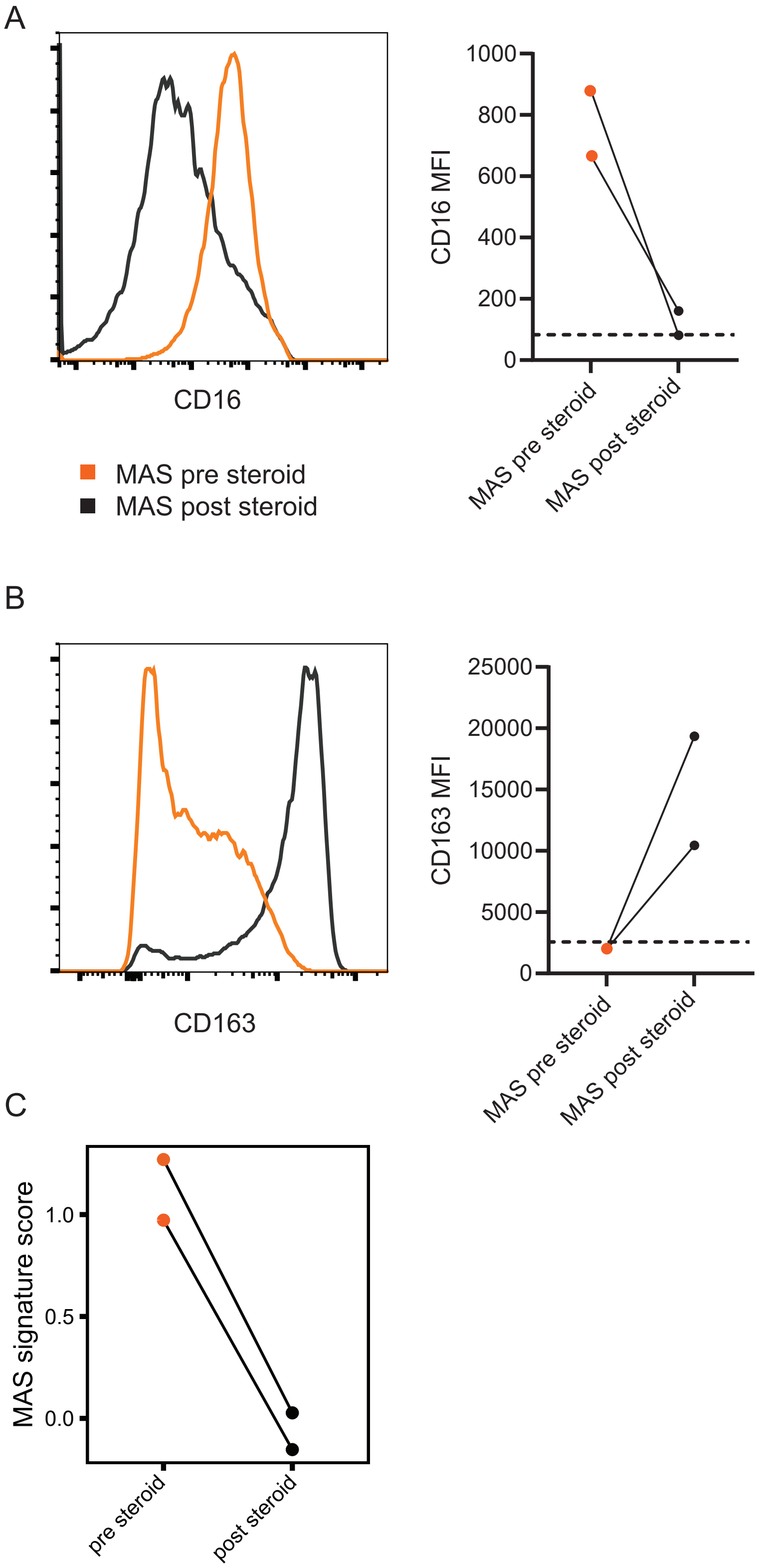

**Supplemental Figure 3: Effect of systemic corticosteroids on monocyte gene expression.** $CD14^+$ monocytes from MAS subjects before (orange) and after (black) systemic corticosteroids were analyzed by flow cytometry for CD16 expression (A) and CD163 (B) expression. On left, histogram shows expression of indicated protein pre (orange) and post (black) steroid treatment for one individual. On right, graph displays MFI for the paired samples from two individuals. Dotted line indicates average MFI for HC. (C) The MAS signature score before (left) and after (right) patients received corticosteroids.

Supplemental Figure 4

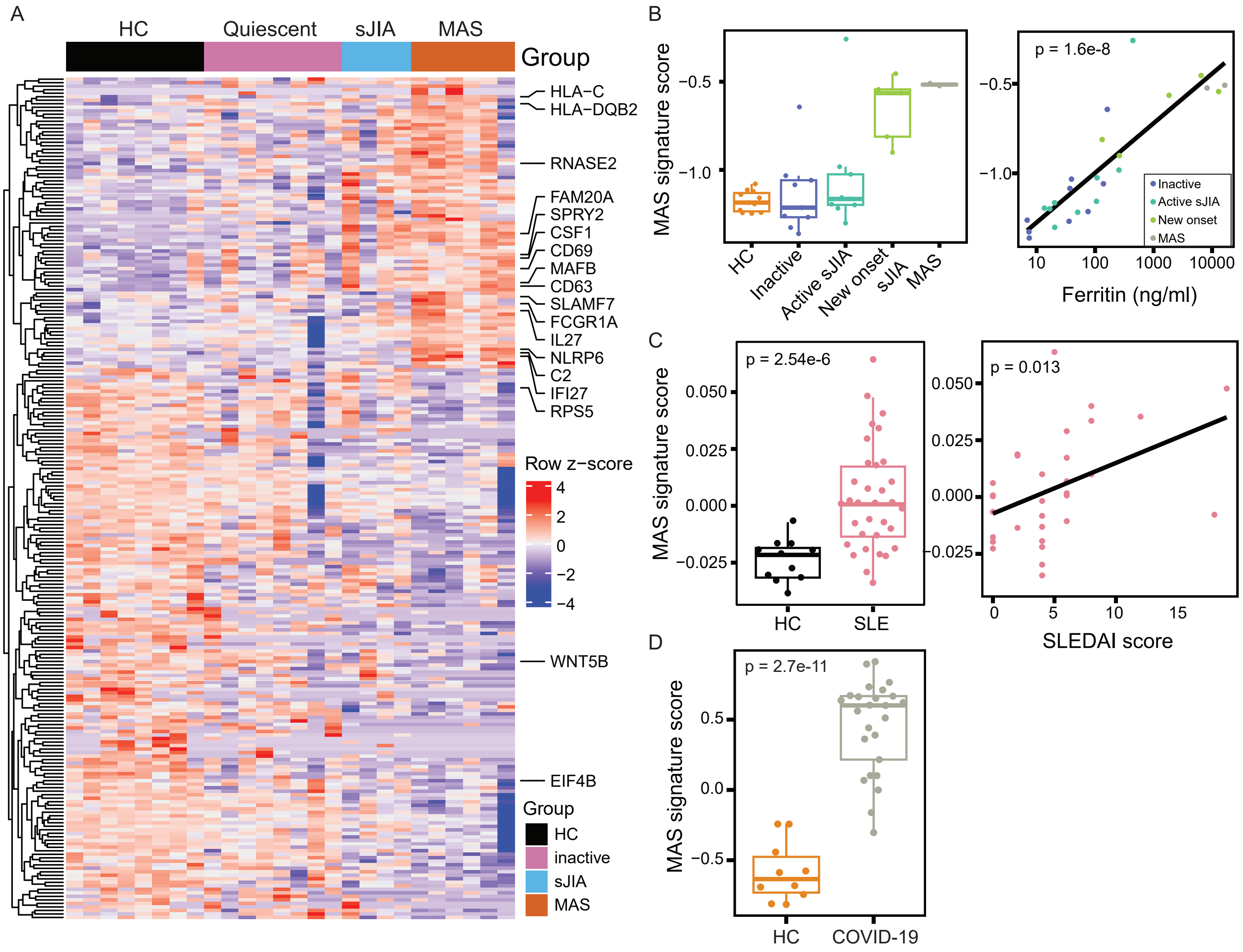

**Supplemental Figure 4: MAS monocyte signature score across data sets.** (A) Heatmap showing DEGs identified between MAS and HC applied to $CD14^{+}CD16^{-/low}$ monocytes from individuals in the following groups: HC (black), quiescent (pink), active sJIA (blue) or active MAS (orange). Z-scores for DEGs are displayed for individual participants (columns). (B) MAS signature score generated in Figure 1C was applied to bulk RNA-Seq data samples from an independent monocyte sJIA dataset from (12) (left). MAS signature score was correlated to ferritin level for the individual samples (right). (C) MAS signature score was applied to pseudobulked scRNA-Seq data of monocytes from pediatric SLE samples from (24) (left). MAS signature score was correlated to SLEDAI (right). (D) MAS signature score was applied to bulk RNA-Seq data from $CD14^{+}CD16^{-/low}$ monocytes sorted from adults hospitalized with COVID-19 or matched HC.

Supplemental Figure 5

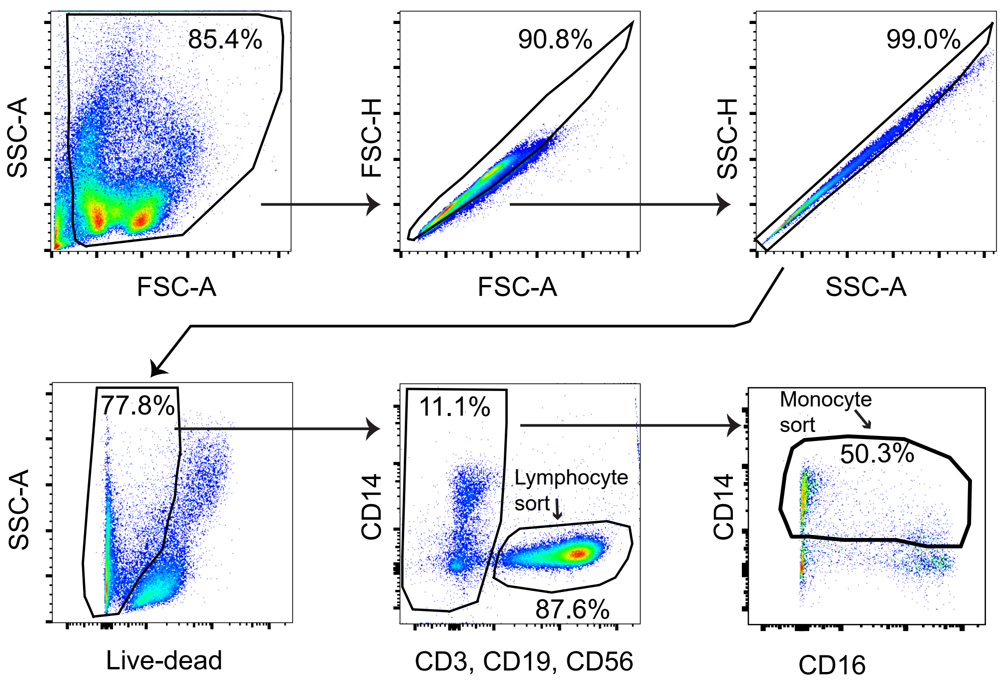

**Supplemental Figure 5: Gating for sorting for monocyte and lymphocyte scRNA-Seq.**

PBMCs were stained with the indicated antibodies and gated as shown to identify monocytes and lymphocytes used in scRNA-Seq analysis.

Supplemental Figure 6

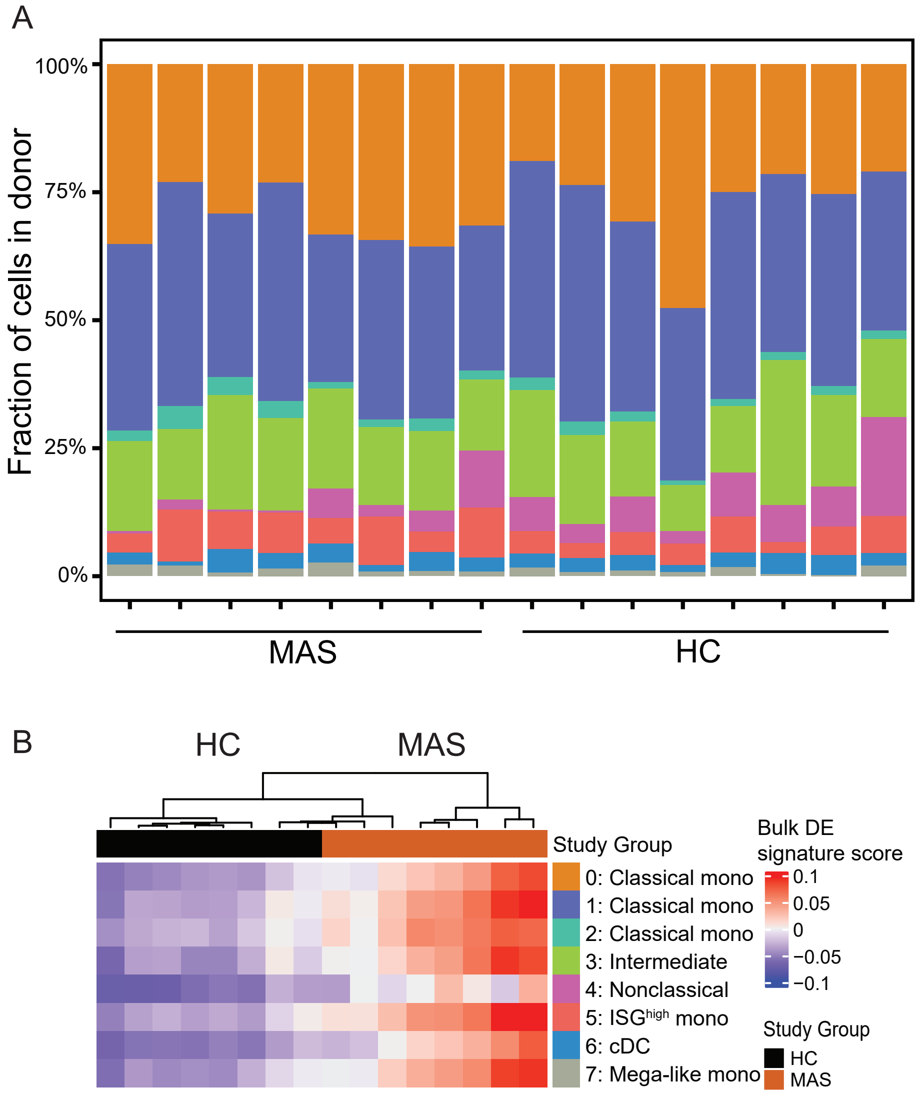

**Supplemental Figure 6: Monocyte proportions and MAS signature score in scRNA-Seq.** (A) Proportion of each cluster from scRNA-Seq analysis of monocytes by individual participant. Clusters are defined in Figure 2B. (B) MAS signature score in scRNA-Seq monocyte clusters. Scaled MAS signatures scores of single cell expression pseudobulked by donor and cluster.

Supplemental Figure 7

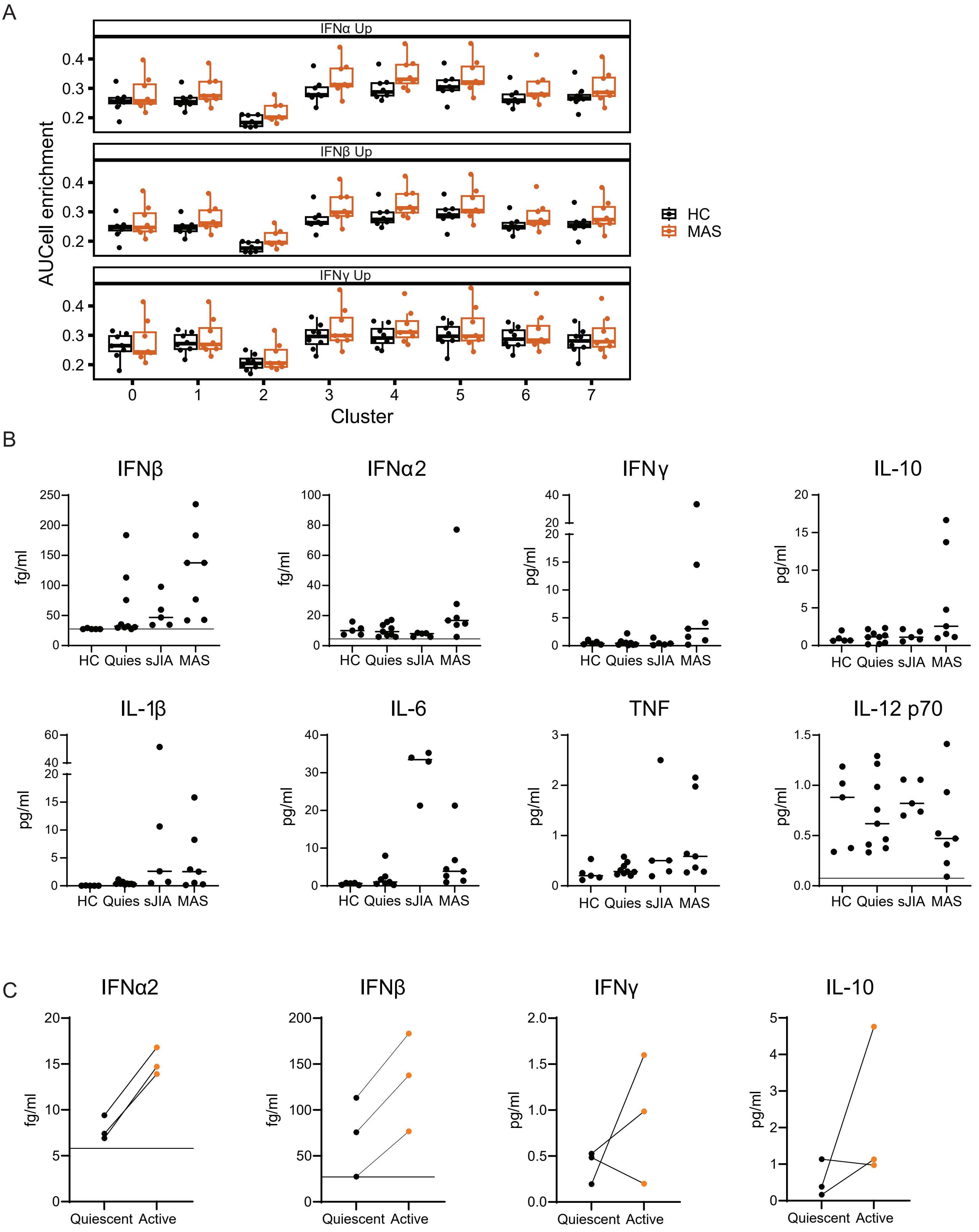

**Supplemental Figure 7: Cytokine expression in sJIA and MAS.** (A) IFN signatures by cluster and individual participant. Area under the curve enrichment for IFN-α2 (top), IFN-β (middle), and IFN-γ (bottom) monocyte signatures from HC (black) or MAS (orange) cells from specific monocyte clusters. Each dot indicates the specific value for an individual participant by cluster and IFN signature. These data gave rise to the calculated values shown in Figure 4D. (B) Plasma concentrations of selected cytokines in HC, quiescent (Quies.), active sJIA, and MAS samples. For those samples below the limit of assay detection, the concentration was set to the level of detection, which is indicated with a horizontal line. For samples below the limit of quantitation but above the limit of detection, the value measured is displayed. Data from individuals on IL-6 inhibiting medications were excluded from IL-6 assessment. IFNα2 and IFNβ are shown as fg/ml. All other cytokines shown as pg/ml. Median for each group is indicated. (C) IFNα2, IFNβ, IFNγ, and IL-10 levels from three individuals with samples collected during quiescent disease and active MAS.

Supplemental Figure 8

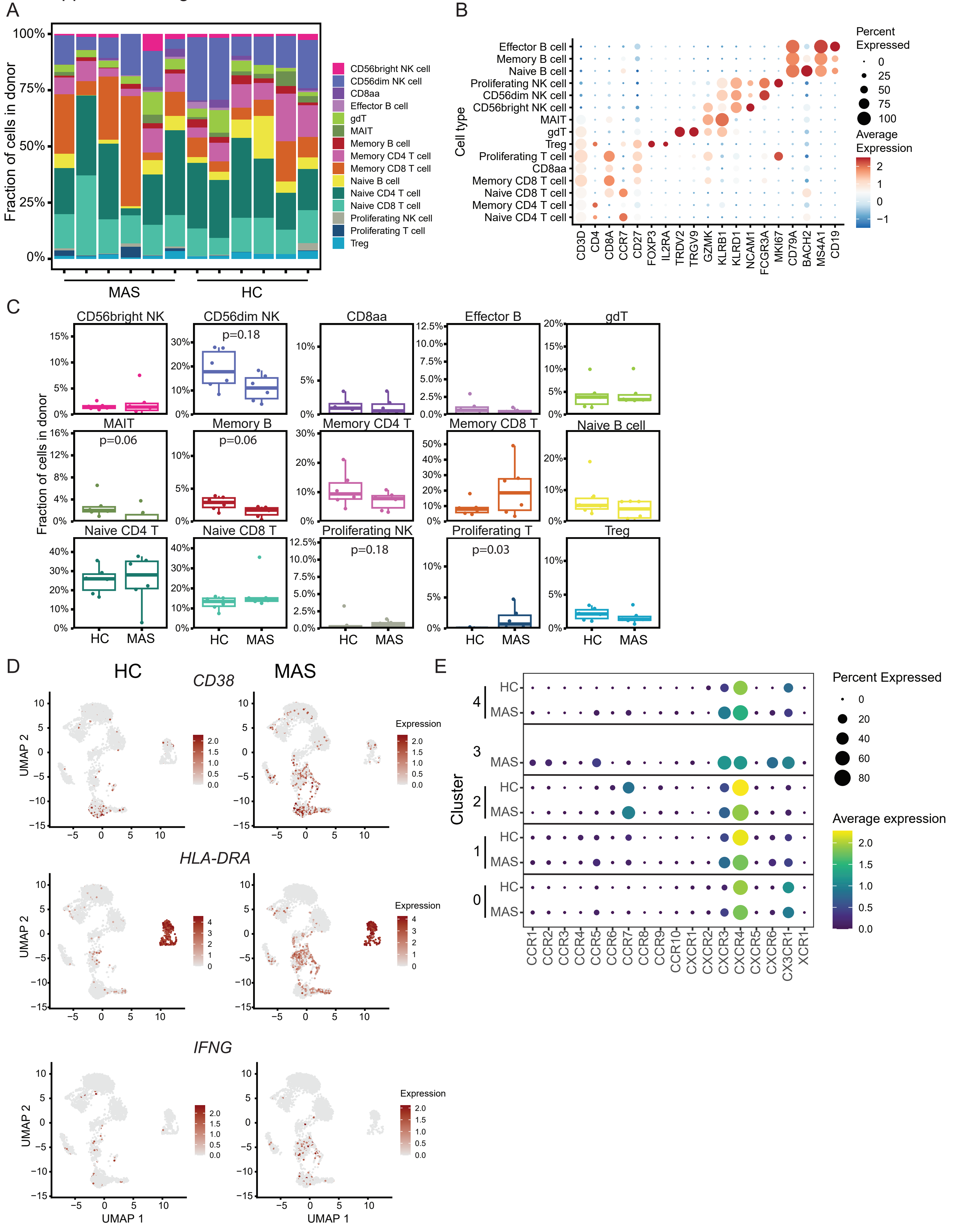

**Supplemental Figure 8: Defining lymphocyte clusters by scRNA-Seq.** (A) Proportion of each cluster from scRNA-Seq analysis of lymphocytes by individual participant. Clusters are defined in Figure 5A. (B) Expression of selected lymphocyte cell type defining genes in lymphocyte clusters shown in Figure 5A. Dot size indicates the percentage of cells expressing the gene, and color reflects the average scaled log-normalized expression. (C) Cell type proportions compared between HC and MAS participants. p-values were calculated using Mann Whitney U test. (D) Expression of *CD38* (top), *HLA-DRA* (middle) and *IFNG* (bottom) expression on HC (left) and MAS (right) lymphocytes, downsampled to 4,000 cells per UMAP. (E) Chemokine receptor expression in effector/memory $CD8^+$ T cell clusters. Size of dot represents percent expressed and color indicates average scaled log-normalized expression.

Supplemental Figure 9

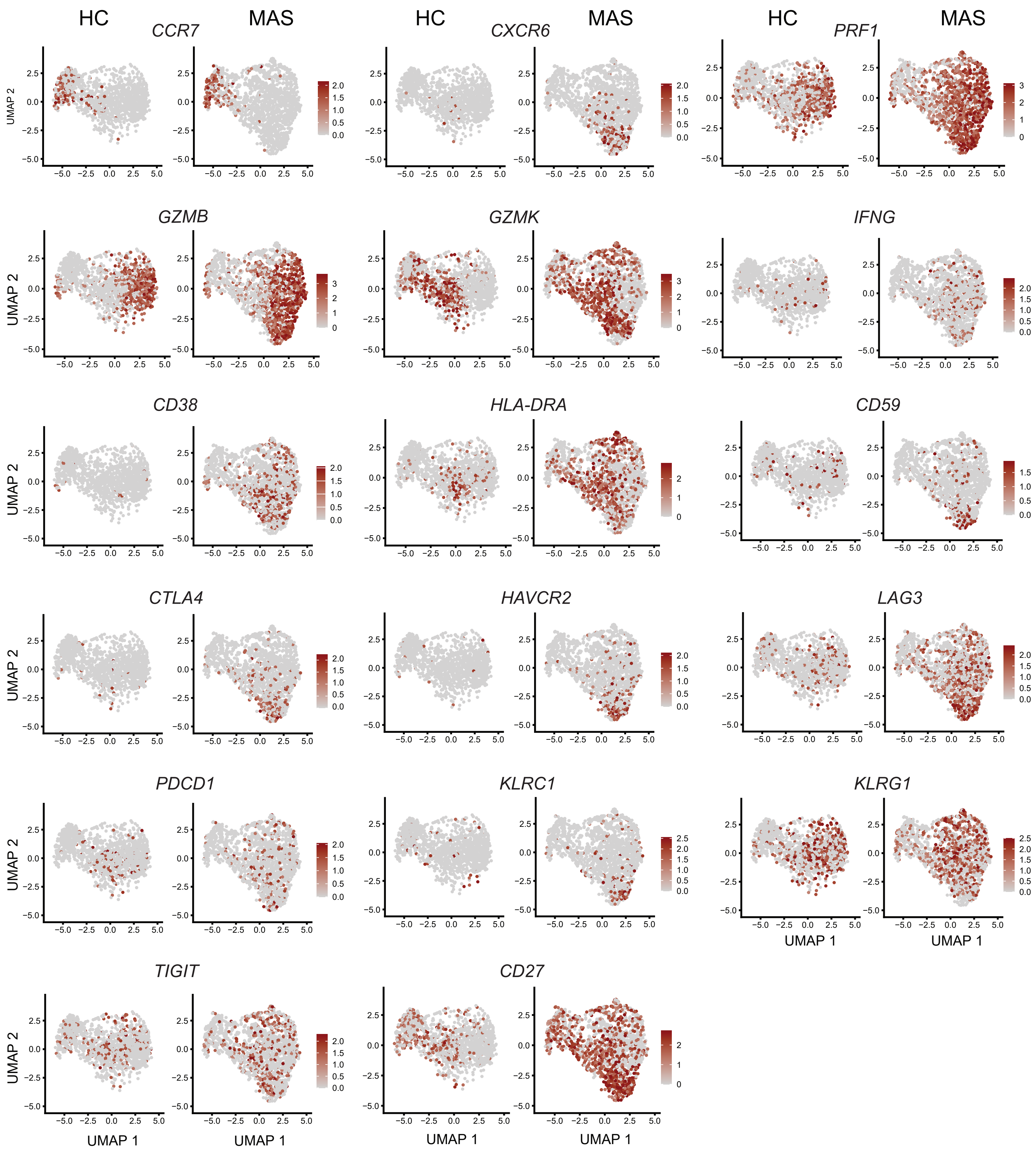

**Supplemental Figure 9: Expression of genes of interest in effector/memory $CD8^+$ T cells.** Specific genes of interest displayed on feature maps. Expression of specific genes displayed as red dots in HC (left) and MAS (right) effector/memory $CD8^+$ T cells, downsampled to 1,500 cells per UMAP.

Supplemental Figure 10

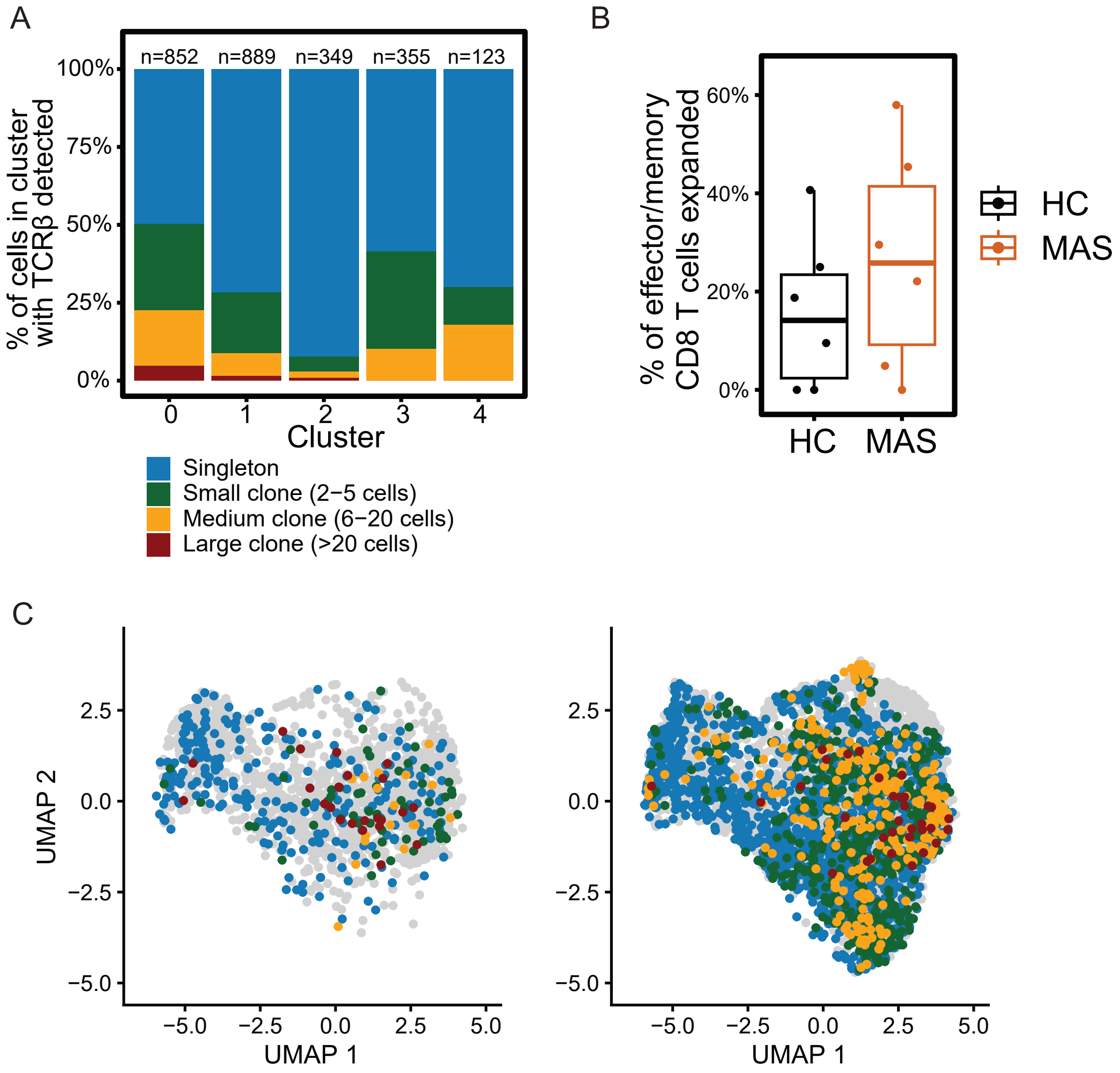

**Supplemental Figure 10: TCR Vβ clonality in effector/memory $CD8^+$ T cells.** (A) Bar plot of TCR Vβ CDR3 clone proportion for each effector/memory $CD8^+$ T cluster. Numbers on top of the bars report the total number of cells where a TCR could be detected for each cluster. Colors indicate number of cells with the same TCR Vβ CDR3 (blue = 1 cell, green = 2-5 cells, yellow = 6-20 cells, red = > 20 cells). Cells where TCRs were not detected were excluded. (B) Percentage of expanded effector/memory $CD8^+$ T cells per donor. Each dot indicates an individual participant. Percentage expanded defined as the number of cells with a TCR Vβ CDR3 clonotype that appears in more than one cell divided by the total number of cells with detected TCR Vβ CDR3. Cells with no TCR Vβ detected are excluded from the plot. $p = 0.38$ by Wilcox test. (C) UMAP for HC (left) or MAS (right) effector/memory $CD8^+$ T cells with color indicating degree of TCR expansion. Gray dots indicate cells in which there was no TCR Vβ CDR3 identified (due to insufficient sequencing depth). For these UMAPs, there was no downsampling and all cells are displayed.

Supplemental Figure 11

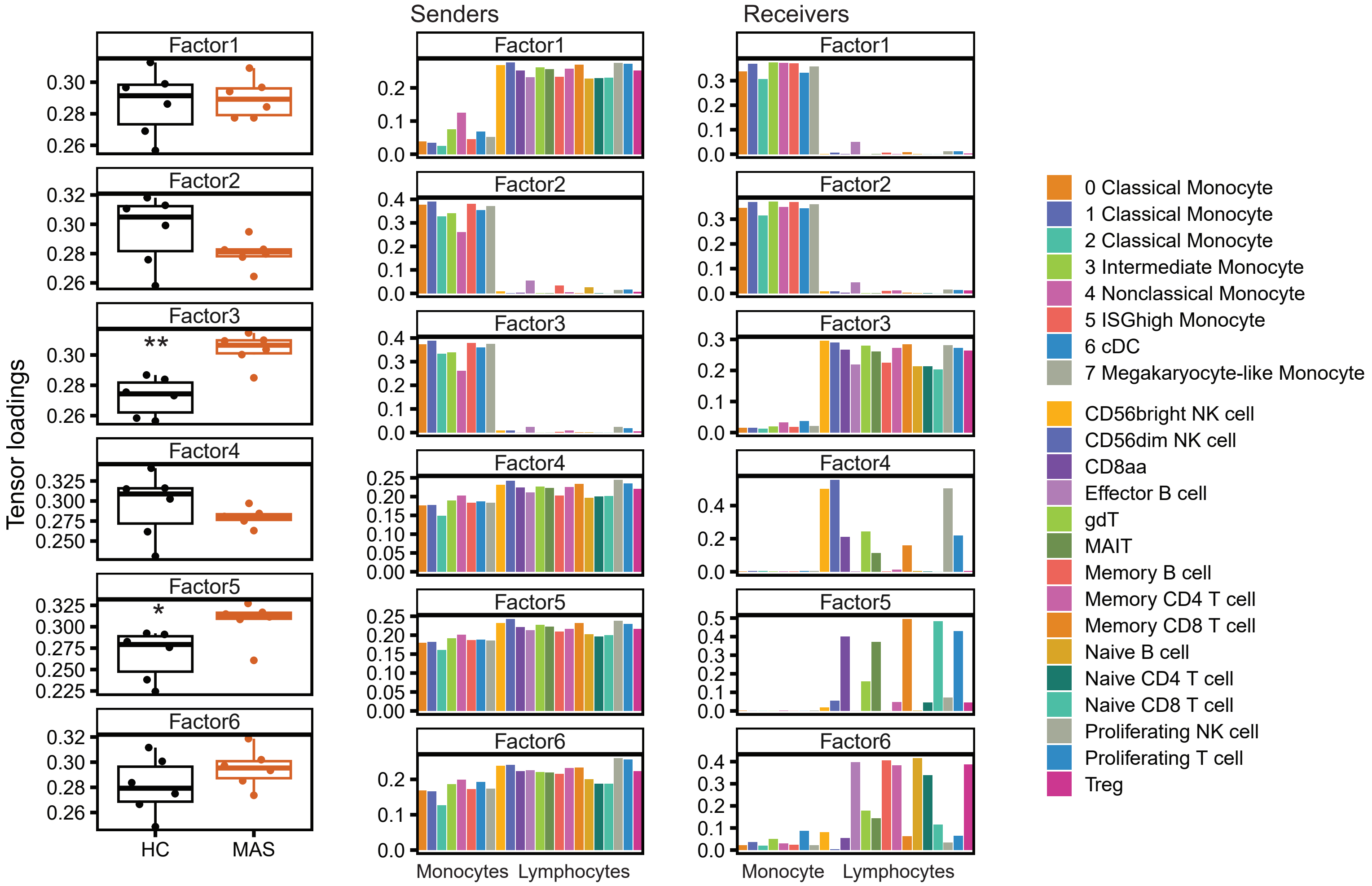

**Supplemental Figure 11: Cell-cell interactions in monocytes and lymphocytes in MAS.**

Tensor-cell2cell data for all six identified factors (left). Only Factors 3 and 5 were significantly different between MAS and HC. Senders (middle) and receivers (left) are shown for each Factor.
